# MetExPred: A Comprehensive Prediction Framework with Protein-Context-Aware Multi-view Learning for Drug Metabolism and Excretion

**DOI:** 10.64898/2026.09.21.753060

**Authors:** Yuxin Huang, Jingting Wan, Gexuyang Wu, Danhong Dong, Yang-Chi-Dung Lin, Hsi-Yuan Huang, Hsien-Da Huang

## Abstract

Within the ADMET continuum, metabolism and excretion (ME) form a critical bridge between drug exposure established by absorption and distribution and downstream efficacy and toxicity. However, no existing framework for drug ME prediction has simultaneously achieved broad coverage of endpoints and robust data recency and completeness. Here, we developed MetExPred, a multi-view prediction framework that comprises 17 classification endpoints and two regression endpoints, covering the overall drug ME process. The framework combines sequence-based and graph-based molecular representations, while optionally incorporating ESM-2 protein representations when experimentally annotated targets are available. A protein-aware masking strategy enables the same architecture to operate in both molecular-only and target-enhanced settings, and multi-view attention adaptively integrates the available representations. Across the classification tasks, MetExPred achieved mean AUROC, AUPRC and F1 scores of 0.844, 0.734 and 0.697, respectively. For clearance and half-life prediction, the model achieved RMSE values of 0.686 and 0.668. MetExPred showed the strongest average performance among the evaluated baselines, while ablation studies confirmed the complementary contributions of molecular sequence, graph and protein information. These results provide a unified and flexible modeling framework for systematic ME prediction of drug compounds, enabling early-stage virtual screening and pharmacokinetic assessment during lead optimization.

**Graphic Abstract:** 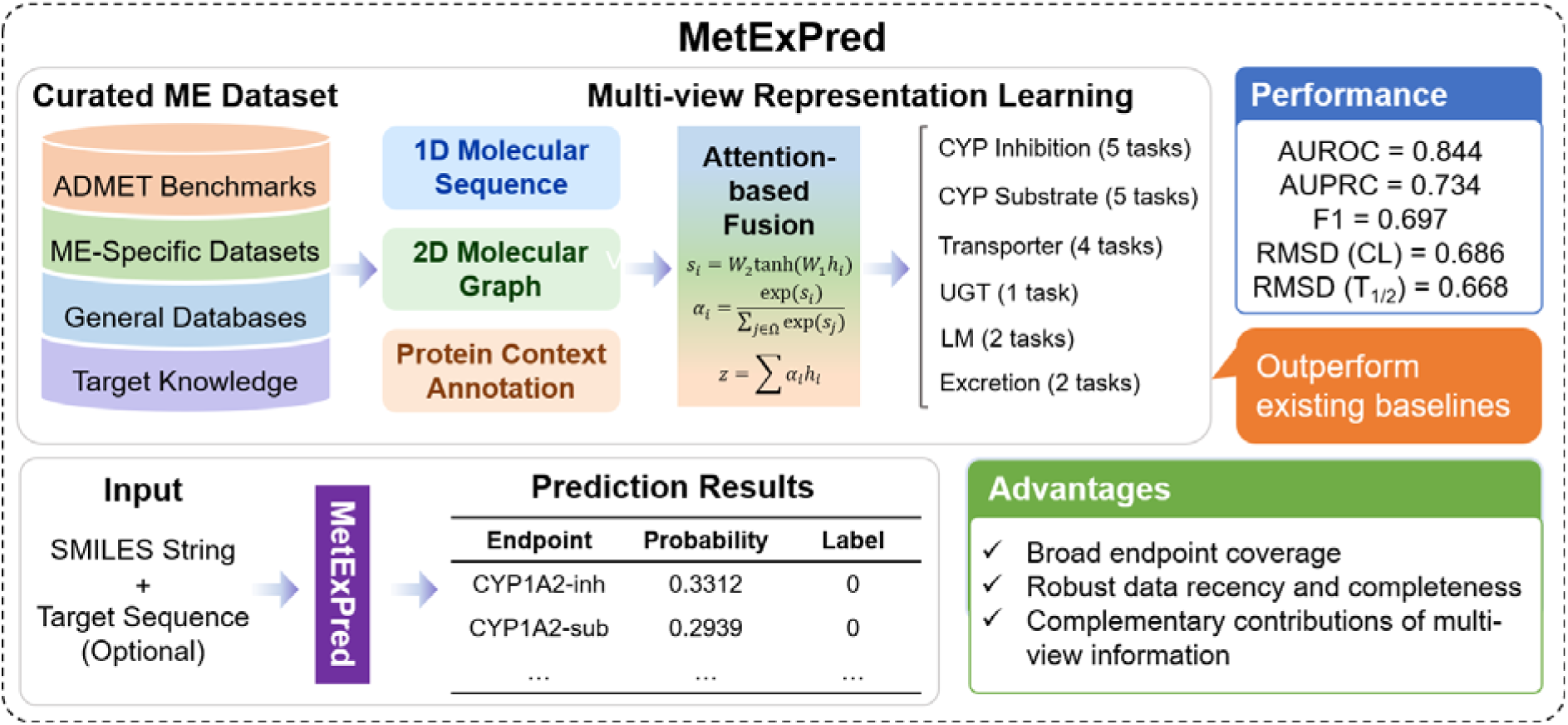

## 1 Introduction

Absorption, distribution, metabolism, excretion and toxicity (ADMET) collectively determine the pharmacokinetic behavior of drug molecules and are fundamental to the efficacy, safety and developability of drug candidates. Early assessment of ADMET properties is therefore essential for identifying unfavorable compounds, reducing development risk and improving decision-making during drug discovery [1–3]. While absorption and distribution determine how a drug enters the systemic circulation and reaches its sites of action, metabolism and excretion (ME) govern its subsequent biotransformation and elimination, thereby linking drug exposure to downstream pharmacological and toxicological outcomes. Metabolic reactions convert parent compounds into metabolites with altered physicochemical and biological properties, whereas excretory processes remove drugs and their metabolites from the body. Together, ME processes regulate systemic elimination, exposure and residence time, and thus play a critical role in preventing excessive accumulation and exposure-related toxicity [4–6]. Drug metabolism is primarily mediated by Phase I and Phase II enzymes. Cytochrome P450 (CYP450) enzymes are major catalysts of Phase I oxidation and related reactions, while uridine diphosphate-glucuronosyltransferases (UGTs) mediate glucuronidation, one of the principal Phase II conjugation pathways [4, 5, 7]. Drug excretion, particularly renal secretion and reabsorption, is strongly influenced by membrane transporters, including OATs, OCTs, MATEs, MRPs, P-glycoprotein and BCRP [8, 9]. Besides commonly-used measurements for drug elimination like plasma clearance and half-life, liver microsomal stabilities are assessed to intuitively reflect clearance rate of chemicals during liver metabolism, acting as a bridge between metabolism and excretion processes [10]. Since metabolic enzymes and transporters jointly determine the persistence and elimination of many drugs, altered ME processes can change systemic exposure, contribute to drug–drug interactions and amplify the risk of therapeutic failure or toxicity. Accordingly, systematic characterization of ME properties provides a critical bridge between early pharmacokinetic behavior and downstream safety assessment, and is also highly relevant to prodrug design and precision medicine, where enzyme-dependent activation and interindividual differences in metabolic or elimination capacity can directly affect treatment outcomes [11, 12].

Experimental evaluation of metabolism and excretion commonly relies on in vitro enzyme or transporter assays, microsomal systems, cellular models and in vivo pharmacokinetic studies. Although these experiments remain indispensable, they are costly, time-consuming and difficult to apply to the large number of compounds generated during early drug discovery. Machine learning (ML) and deep learning (DL) provide a scalable alternative by learning relationships between molecular information and experimentally measured ADMET properties, thereby enabling rapid prioritization before extensive experimental validation. Early ADMET prediction methods mainly used molecular descriptors or fingerprints with conventional algorithms such as random forests, support vector machines and other quantitative structure-activity relationship models. Representative resources including admetSAR, pkCSM, SwissADME, ADMETlab and admetSAR 2.0 demonstrated the feasibility of broad in silico assessment across multiple pharmacokinetic and toxicity endpoints [13–17]. With the expansion of curated chemical datasets and advances in molecular representation learning, recent ADMET prediction has increasingly shifted toward graph neural networks, message-passing architectures and multi-task learning. Platforms such as ADMETlab 2.0, Interpretable-ADMET, ADMETboost, ADMETlab 3.0, Deep-PK and admetSAR 3.0 have substantially expanded endpoint coverage and adopted more advanced ML or DL strategies for large-scale prediction [18–23]. In particular, current deep models can learn task-relevant features directly from molecular graphs and share information across related endpoints, reducing dependence on manually designed descriptors. These developments have greatly improved general ADMET prediction; however, broad ADMET platforms are designed to cover heterogeneous pharmacokinetic and toxicity properties, and metabolism- and excretion-related endpoints usually represent only a subset of their prediction space. Consequently, the biological relationships among metabolic enzymes, transporters, substrates and inhibitors are not usually the central focus of model or dataset design.

ME-specific prediction has developed mainly around CYP enzymes, UGT-mediated metabolism and individual drug transporters, with CYP-related tasks receiving the greatest attention. Early CYP studies commonly focused on a specific enzyme or narrowly defined prediction task. RS-Predictor, for example, predicted sites of CYP3A4-mediated metabolism, whereas proteochemometric modeling was used to predict inhibition across multiple CYP isoforms by jointly representing compounds and proteins [24, 25]. Subsequent work expanded CYP prediction toward reactant and inhibitor classification. CypReact predicts whether a molecule is a reactant for nine major human CYP enzymes and uses cost-sensitive learning to reduce the impact of false-negative reactant predictions [26]. SuperCYPsPred and CYPlebrity developed isoform-specific CYP inhibition models using larger datasets and sampling or balancing strategies to address the strong imbalance between inhibitors and non-inhibitors [27, 28], while consensus QSAR approaches have also been explored for CYP inhibition and induction [29]. More recently, DL has enabled multi-task and graph-based modeling of CYP activity. DEEPCYPs combines fingerprint and graph representations for multi-task prediction of five major CYP isoforms [30]. The scope of computational metabolism has also extended beyond binary substrate or inhibitor prediction. Deep learning has been applied to metabolite prediction [31], and recent frameworks such as DeepMetab and DeepCYP integrate multiple stages of CYP-mediated metabolism, including substrate profiling, site-of-metabolism localization and metabolite or product prediction [32, 33]. Compared with CYP prediction, computational modeling of UGT-mediated metabolism and transporter-related excretion remains less extensive. UGT models have largely focused on glucuronidation site prediction [34, 35] or inhibition of selected enzymes such as UGT1A1 [36]. Transporter models are often developed for a single protein; for example, OATP1B1 inhibition models have been built from experimentally generated transport data and used to identify previously unreported inhibitors [37]. Although these studies have advanced individual ME tasks, the overall field remains fragmented. CYP isoforms, UGT enzymes and transporters are typically modeled in separate studies with independently collected datasets, different endpoint definitions and task-specific architectures. As a result, existing models provide strong solutions for selected metabolic reactions or proteins but do not yet offer a unified framework for systematically characterizing a drug across the broader metabolism-excretion process.

Taken together, these studies also reveal a common limitation in current computational ADMET and ME modeling. Representative broad ADMET platforms and recent ME-specific deep-learning methods formulate predictions primarily from molecular descriptors, fingerprints, SMILES, or molecular graphs [21–23, 30, 32]. Although proteochemometric modeling has jointly represented compounds and CYP isoforms [25], the protein representation in that setting defines the enzyme-specific prediction task, which differs from using a compound’s broader pharmacological target profile as an auxiliary source of biological context. Across these representative methods, such compound-associated target profiles have not been systematically evaluated across a harmonized set of heterogeneous ME endpoints [21–23, 25, 30, 32]. Nevertheless, intended targets and experimentally measured compound-target activities are generated during hit identification and lead optimization and may therefore already be available for at least a subset of prioritized compounds before clinical pharmacokinetic outcomes are established [38, 39]. This creates an opportunity to investigate whether target information can complement molecular representations in systematic ME prediction.

To address these limitations, we constructed a comprehensive dataset comprising 17 metabolism-related classification endpoints and two excretion-related regression endpoints, covering major CYP isoforms, UGT-mediated metabolism, drug transporters, liver microsomal stabilities, clearance and half-life. Based on the preprocessed data, we developed MetExPred, a target-aware multi-view framework that integrates 1D molecular sequence representations derived from ChemBERTa, 2D molecular graph representations learned by a directed message-passing neural network (D-MPNN), and ESM-2 protein representations from experimentally annotated compound-associated targets. These views provide complementary information on molecular semantics, structural topology and biological context. A protein-aware masking strategy handles missing target annotations, while a multi-view attention module adaptively integrates the available representations for each prediction task. The resulting architecture supports both classification and regression endpoints within a unified framework, enabling systematic prediction of macro rate measurements and micro drug-enzyme interactions. By combining complementary molecular and target information, MetExPred provides an integrated approach for early pharmacokinetic assessment and identification of metabolism- and transporter-related liabilities.

## 2 Materials and Methods

### 2.1 Dataset construction and preprocessing

We constructed an ME-oriented dataset containing 19 endpoints: 17 metabolism-related classification tasks and 2 excretion-related regression tasks. The classification endpoints comprised inhibition and substrate activities for CYP cytochromes CYP1A2, CYP2C19, CYP2C9, CYP2D6 and CYP3A4; inhibitor activities for transporters OATP1B1, OATP1B3, OCT1 and OCT2; UGT substrate activity; and human liver microsomal (HLM) and rat liver microsomal (RLM) stability. Plasma clearance (shortened as clearance) and half-life were included as continuous excretion-related outcomes with a log transformation. Together, these endpoints represent enzyme-mediated biotransformation, transporter-associated disposition, metabolic stability and systemic elimination.

Data were integrated from three complementary source categories (Figure 1A). First, ML-ready benchmark data from Deep-PK [22] and admetSAR 3.0 [23] were incorporated to retain compatibility with established ADMET evaluation settings. Second, specialized resources were used to extend CYP interaction, transporter inhibitor/substrate, microsomal stability, clearance and half-life annotations [40–46]. Third, DrugBank [47] and ChEMBL 36 [48] were used to expand chemical coverage and supplement compound and bioactivity annotations. All records were subjected to a unified preprocessing pipeline comprising structure validation, canonical simplified molecular input line entry system (SMILES) generation, salt removal, duplicate handling and mapping to consistent endpoint definitions and label formats.

**Figure 1.**
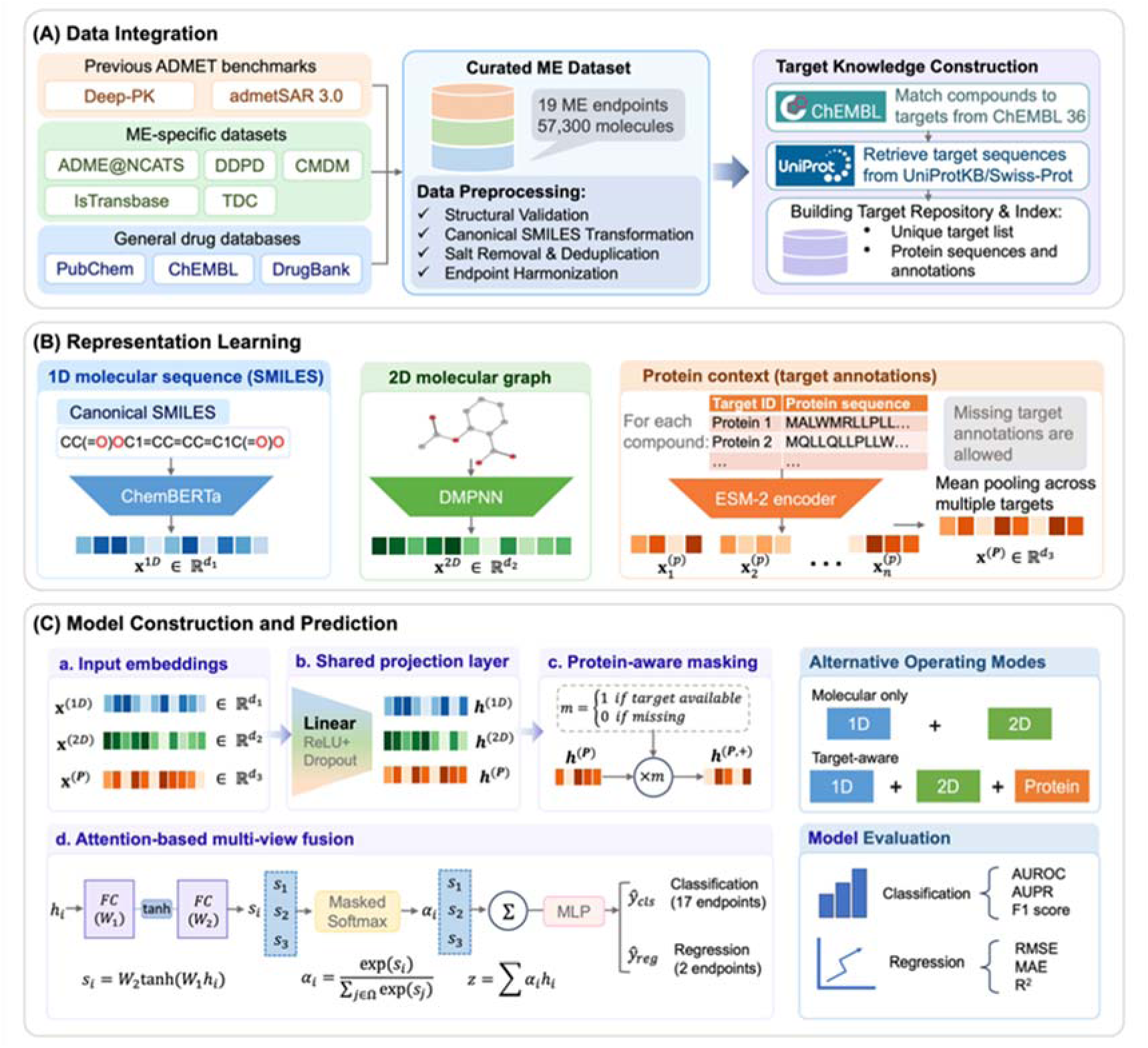
Overview of the MetExPred workflow and model architecture. (A) Integration and harmonization of metabolism and excretion data from ADMET benchmarks, ME-specific datasets, and general drug databases, together with construction of a ChEMBL- and UniProt-derived target knowledge repository. (B) Multi-view representation learning using ChemBERTa for 1D molecular sequences, D-MPNN for 2D molecular graphs, and ESM-2 for available target protein sequences, with mean pooling used to obtain compound-level protein representations. (C) Protein-aware multi-view prediction framework. Modality-specific embeddings are projected into a shared latent space, masked when protein annotations are unavailable, and integrated by attention-based fusion before task-specific prediction. The framework supports both molecular-only and target-aware modes and produces outputs for 17 classification and two regression endpoints.

Detailed endpoint definitions, source mapping and sample counts before and after deduplication are reported in Table 1. For each endpoint, compounds were divided into training, validation, and test subsets at a ratio of 8:1:1 using a random split (Table S1). Stratified sampling was applied to classification tasks to preserve endpoint-specific class distributions.

**Table 1.**
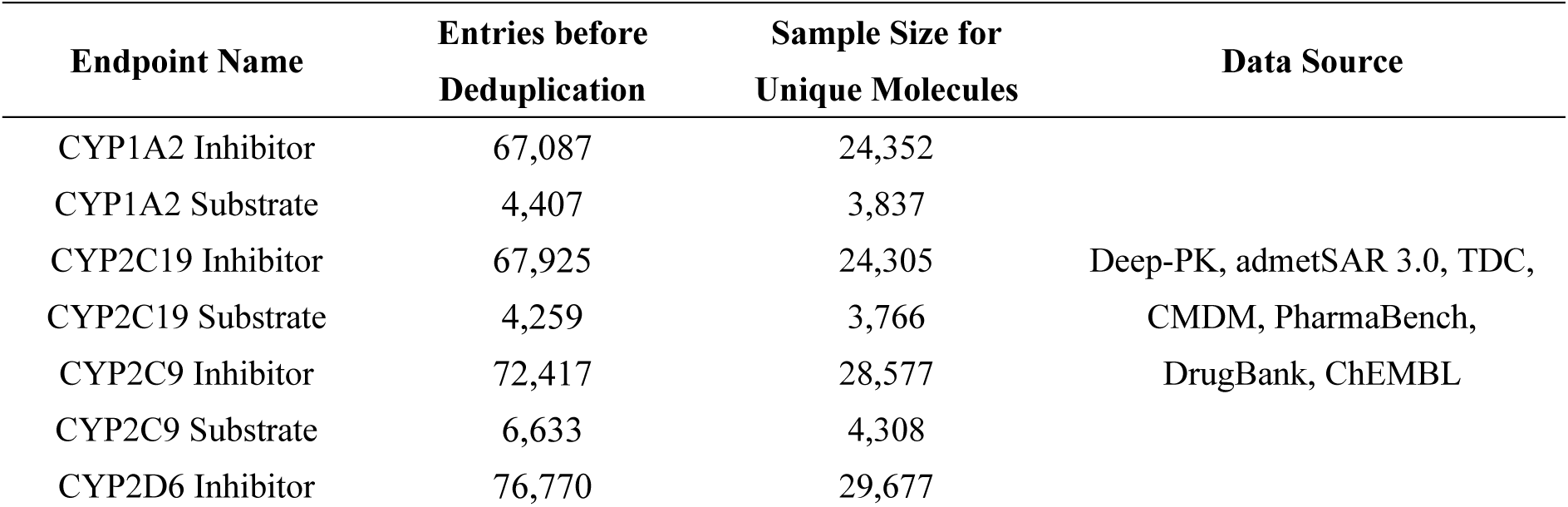
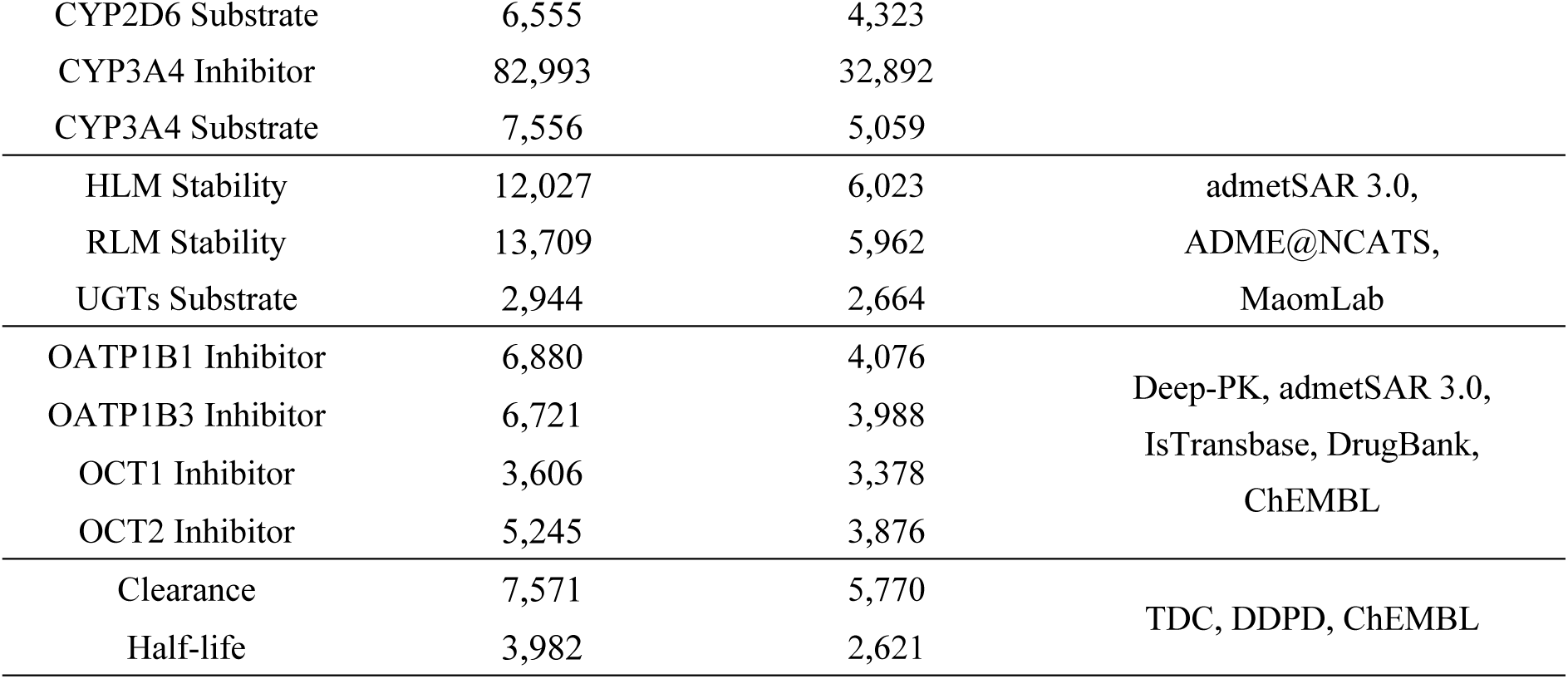
Endpoint name, total sample size and data source for integrated datasets.

### 2.2 Target knowledge construction

To incorporate biological information related to target genes and functional annotations beyond molecular structure, experimentally reported compound-associated protein targets were retrieved from ChEMBL 36 [48]. Compounds in the integrated ME dataset were matched to their reported targets, and the corresponding reviewed amino acid sequences were obtained from UniProtKB/Swiss-Prot using UniProt accessions [49]. One-to-many compound-target relationships were retained because individual compounds may be associated with multiple proteins. The statistics of target annotation coverage across different ME endpoints, including the number of unique targets, target-annotated molecules and molecule–target associations are summarized in Table S2. The resulting target-annotated subset was used for experiments that explicitly incorporated protein representations, whereas compounds without validated target annotations remained available for molecular-only modeling.

All unique proteins were consolidated into a nonredundant target repository, and a target index was created to map each molecular sample to its associated protein sequences. For compounds associated with multiple targets, individual protein embeddings were aggregated by mean pooling to produce one compound-level target representation. This construction provides a general biological activity view; it does not assume that every annotated protein directly mediates the corresponding ME endpoint.

### 2.3 Multi-view representation learning

MetExPred represents each compound through three complementary views (Figure 1B). The one-dimensional (1D) view encodes canonical SMILES using the pretrained ChemBERTa-77M-MTR molecular language model [50]. The two-dimensional (2D) view represents the molecular graph using a D-MPNN, which propagates information along directed bonds to capture atom-bond topology and local chemical environments [51]. For the target-aware subset, protein sequences are encoded using the pretrained ESM2-t33-650M-UR50D model [52]. Each encoder produces a modality-specific dense vector before cross-view integration.

Given the 1D molecular sequence *S_i_*, the molecular graph *G_i_* and the associated target set *T_i_* = {*t_1_*, *t_2_*, …, *t_N_*} for compound *i*, the view-specific embeddings are defined as follows:

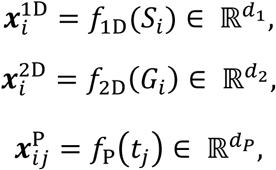

where *f*_1D_, *f*_2D_ and *f*_P_ denote the 1D molecular, 2D molecular and protein embedding functions, respectively.

For compounds with *N* multiple annotated targets, the individual protein embeddings are aggregated by mean pooling:

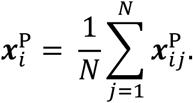

Each compound is consequently represented by a set of available modality-specific embeddings:

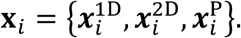

Before fusion, all modality-specific embeddings are projected into a shared latent space of dimension *d*:

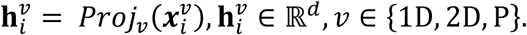

### 2.4 Protein-aware representation masking

Target annotations were unavailable for a proportion of compounds. Directly padding missing protein embeddings and including them in attention computation could introduce artificial signals. We therefore defined a binary protein mask *m_i_*, where *m_i_* = 1 denotes a valid target representation and *m_i_* = 0 denotes missing target information. The projected protein embedding was masked before multi-view fusion:

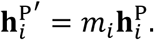

This operation suppresses unavailable protein representations and restricts the attention mechanism to informative views. Molecular-only samples can therefore be processed without treating a padded target vector as biological evidence.

### 2.5 Attention-based view fusion

The relative importance of sequence, graph, and target information may vary across compounds and endpoints. MetExPred therefore learns a view-specific attention score for each available projected representation. The scoring network consists of two fully connected layers with a hyperbolic tangent activation:

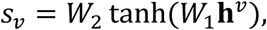

where *W_1_* and *W_2_* are trainable parameters.

The unconstrained scores are normalized across available views using the softmax function:

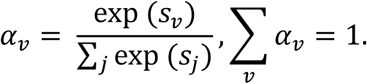

The final fused representation is calculated as a weighted sum of the projected view embeddings with the normalized attention weights:

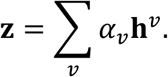

This design allows informative views to contribute more strongly to the final compound representation while retaining complementary chemical and biological information. Figure 1C summarizes the complete workflow from data integration and target annotation to multi-view encoding, masked attention fusion, and endpoint-specific prediction.

### 2.6 Prediction heads, optimization, and evaluation

The fused representation is passed to a multilayer perceptron (MLP) with three fully connected layers and nonlinear activations. A task-dependent output layer is used for binary classification or regression. Classification outputs are converted to probabilities using a sigmoid activation, whereas regression outputs are generated without additional activation.

All trainable components are optimized jointly by backpropagation. Binary classification tasks use binary cross-entropy loss, and regression tasks use smooth L1 loss. The Adam optimizer, dropout regularization, validation-based early stopping, and checkpoint selection are applied during training. Classification performance is evaluated using the area under the receiver operating characteristic curve (AUROC), area under the precision-recall curve (AUPRC), and F1 score. The decision threshold for each endpoint is selected on the validation set by maximizing the F1 score over thresholds from 0.05 to 0.95 and is then fixed for test-set evaluation. Regression performance is measured using root mean squared error (RMSE), mean absolute error (MAE), and the coefficient of determination (R-squared).

### 2.7 Representation, fusion, baseline, and ablation experiments

A controlled representation benchmark was first performed to identify suitable 1D and 2D encoders. Term frequency-inverse document frequency (TF-IDF) [53], ChemBERTa [50], and MoLFormer [54] were evaluated as sequence-based representations, while Morgan fingerprints and D-MPNN embeddings were evaluated as structure-based representations. Individual views and directly concatenated 1D+2D combinations were compared using the same MLP prediction head. The selected ChemBERTa and D-MPNN representations were then evaluated under four fusion strategies: direct concatenation, linear attention, convolutional neural network-based attention, and the proposed multi-view attention module.

For external method comparison, MetExPred was evaluated against baseline methods Chemprop [51], SuperCYPsPred [27], Deep-PK [22], DeepMetab [32], and ADMETlab 3.0 [21]. All methods were trained and evaluated on the same curated endpoint data, splits and metrics. When official implementations or compatible pretrained models were unavailable, architectures were re-implemented according to published descriptions and tuned using the common validation protocol.

Ablation experiments were conducted in two settings. On the complete molecular dataset, 1D-only, 2D-only, and 1D+2D configurations were compared. On the target-annotated subset, 1D+target, 2D+target, 1D+2D, and 1D+2D+target configurations were evaluated to quantify the incremental contribution of protein information.

### 2.8 CYP inhibition case-study design

To illustrate the practical application of MetExPred from complementary perspectives, two sets of case studies were conducted. First, five clinically relevant compounds with diverse CYP inhibition profiles and scaffold characteristics were selected from DrugBank [47]: encorafenib, methylene blue, relacorilant, sertraline, and adagrasib. Molecular-only and target-aware predictions were compared across five CYP inhibitor endpoints. To examine the relationship between prediction behavior and structural familiarity, each compound was further compared with the training set using maximum common substructure-based matching and Tanimoto similarity. Prediction scores, endpoint-specific decision thresholds, DrugBank annotations, and structural similarity were considered jointly.

Separately, a broader set of representative drugs with experimentally or clinically reported metabolism and excretion properties was curated from the literature [55–59]. These literature-grounded cases covered CYP substrate and inhibitor activity, UGT substrate activity, transporter inhibition, microsomal stability, clearance, and half-life. For classification endpoints, literature-supported positive or negative annotations were compared with model prediction probabilities and predicted classes. For regression endpoints, reported pharmacokinetic values were compared with model predictions after back-transformation from the log scale. These cases were used to assess the consistency of MetExPred predictions with reported ME properties across diverse endpoint types, rather than as an independent benchmark dataset.

## 3 Results

### 3.1 Construction and characterization of the ME benchmark

The final benchmark covered 17 metabolism-related classification endpoints and 2 excretion-related regression endpoints. Relative to existing ML-ready benchmarks, the integrated dataset increased the sample size of all shared CYP-related tasks and added several endpoints that were absent or sparsely represented in the comparison resources. In total, the benchmark covers 199,454 data entries and 57,300 unique molecular structures, with an overall scaffold ratio of 0.4991, indicating high structure diversity. The largest classification datasets contained more than 28,000 compounds for CYP2C9, CYP2D6, and CYP3A4 inhibition, while the smaller substrate and transporter tasks retained several thousand compounds. The two regression datasets contained 5,770 clearance records and 2,621 half-life records, surpassing all other sources involved in the data scale comparison (Figure 2A).

**Figure 2.**
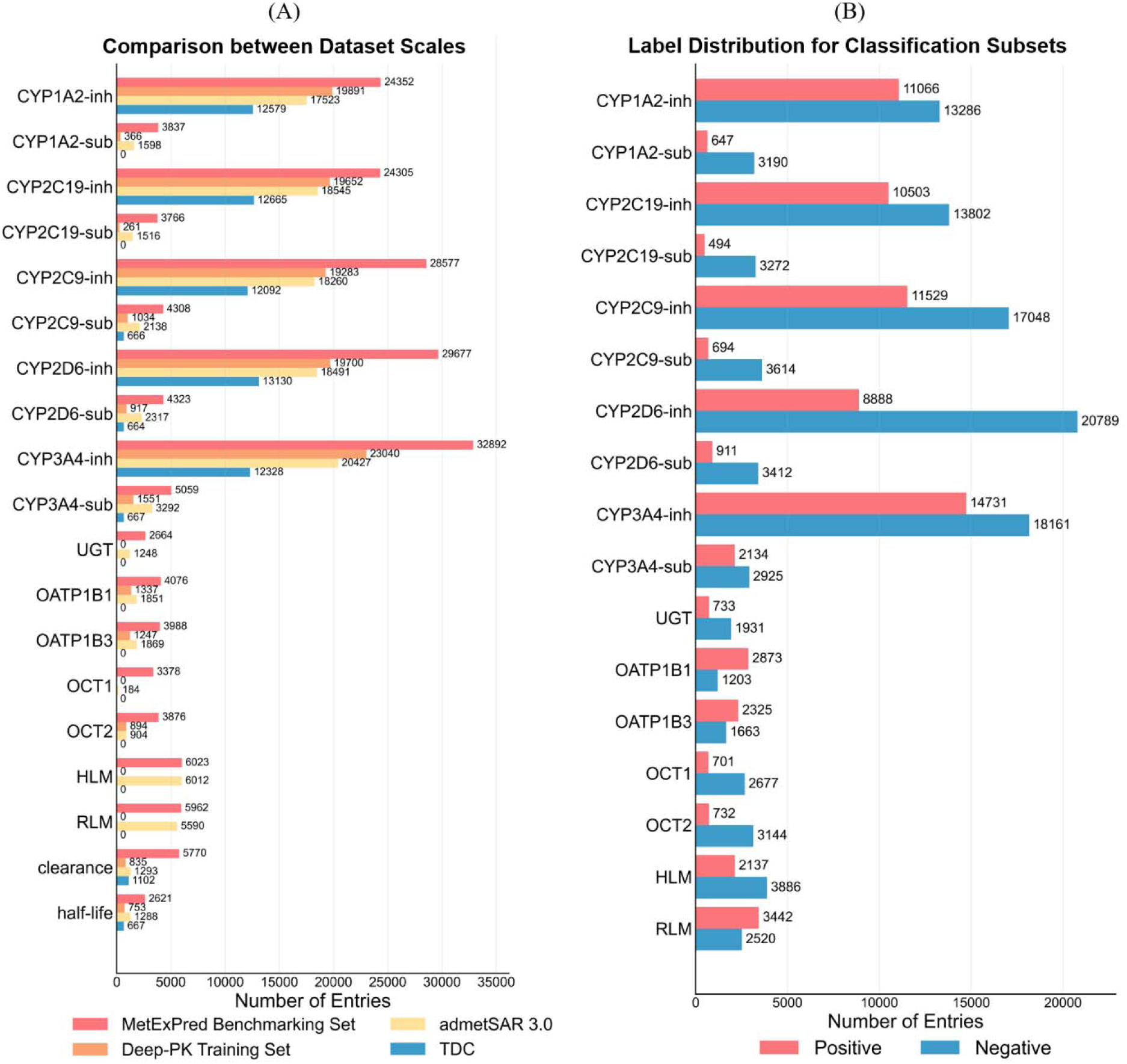
Composition of the curated MetExPred benchmark. (A) Comparison of dataset scales with existing benchmarks for shared metabolism and excretion endpoints. (B) Positive and negative label distributions for the 17 classification endpoints of MetExPred.

The classification endpoints exhibited task-dependent imbalance, reflecting differences in the availability of experimentally validated positive and negative records. Nevertheless, the unified preprocessing and stratified splitting procedures retained both classes in each subset and enabled consistent model evaluation across heterogeneous ME tasks (Figure 2B).

Experimentally reported targets were available for every endpoint, although coverage varied across tasks from approximately 5% to more than 40% of compounds (Figure 3A). The mean number of annotated targets per target-associated compound also differed among endpoints, demonstrating that the target-specific subset retained one-to-many compound-protein relationships rather than reducing each compound to a single annotation (Figure 3B). Functional characterization showed that the target repository included kinases, enzymes, G protein-coupled receptors, transcription factors, ion channels, transporters, and other protein families (Figure 3C). This diversity supports the use of protein representations as a broad biological context layer but also emphasizes that the annotations are not restricted to direct metabolic enzymes or excretion transporters.

**Figure 3.**
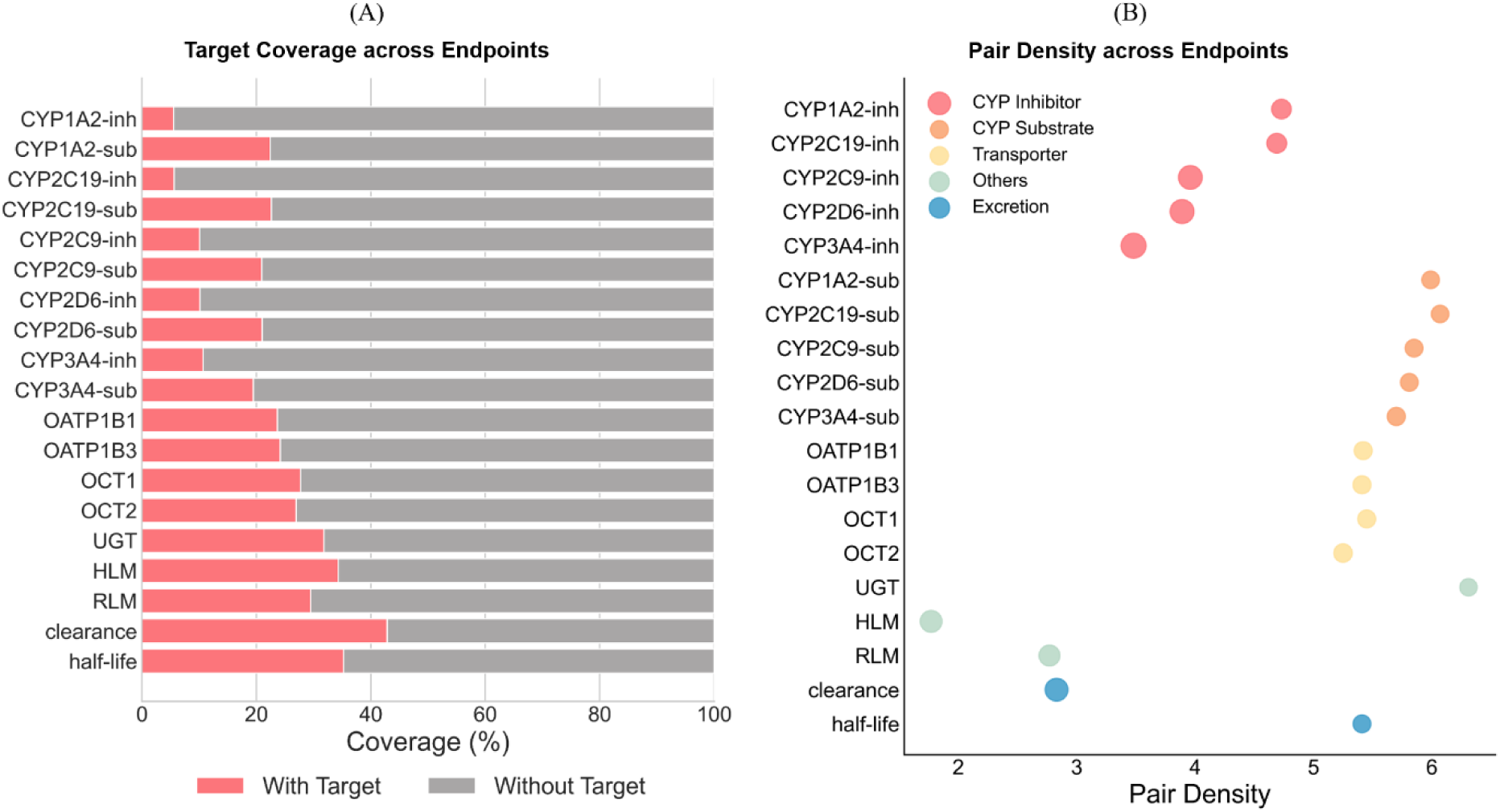
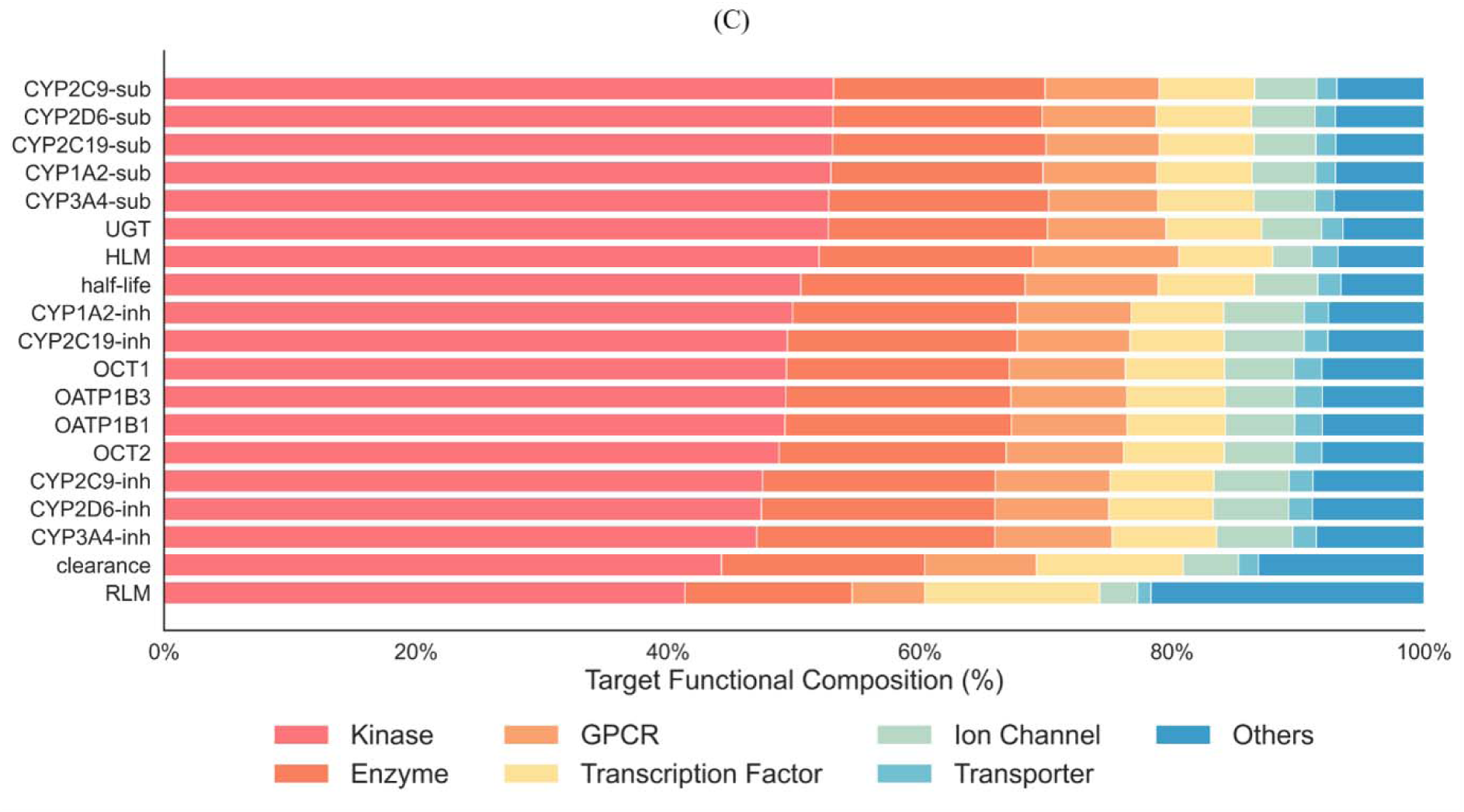
Construction and characterization of the target-aware subset. (A) Proportion of compounds with and without experimentally annotated targets across endpoints. (B) Mean compound-target pair density for each endpoint. (C) Functional composition of the annotated protein targets.

### 3.2 Overall performance compared with baseline methods

Preliminary component comparisons identified ChemBERTa and D-MPNN as the most effective 1D and 2D molecular encoders, respectively, and showed that their combination consistently outperformed either representation alone (Figure S1 and Table S3-S6). Among the tested fusion mechanisms, multi-view attention provided the strongest aggregate performance, supporting its use in the final MetExPred architecture (Table S7-S10). We therefore placed the principal benchmark comparison before the detailed design and ablation analysis.

Under the full-set molecular configuration, MetExPred achieved mean AUROC, AUPRC, and F1 values of 0.8444, 0.7340, and 0.6966 across the 17 classification endpoints. The model also achieved an RMSE of 0.6856, an MAE of 0.4469, and an R-squared value of 0.5000 for clearance, as well as an RMSE of 0.6683, an MAE of 0.4735, and an R-squared value of 0.2494 for half-life (Table 2). Across the common benchmark, MetExPred showed the strongest average performance among the evaluated ADMET and metabolism-specific baselines, while individual methods remained competitive on selected endpoints (Figure 4A). This endpoint-level variation is expected because part of ME tasks are driven primarily by local chemical substructures, whereas others depend on broader molecular context or biological interactions.

**Figure 4.**
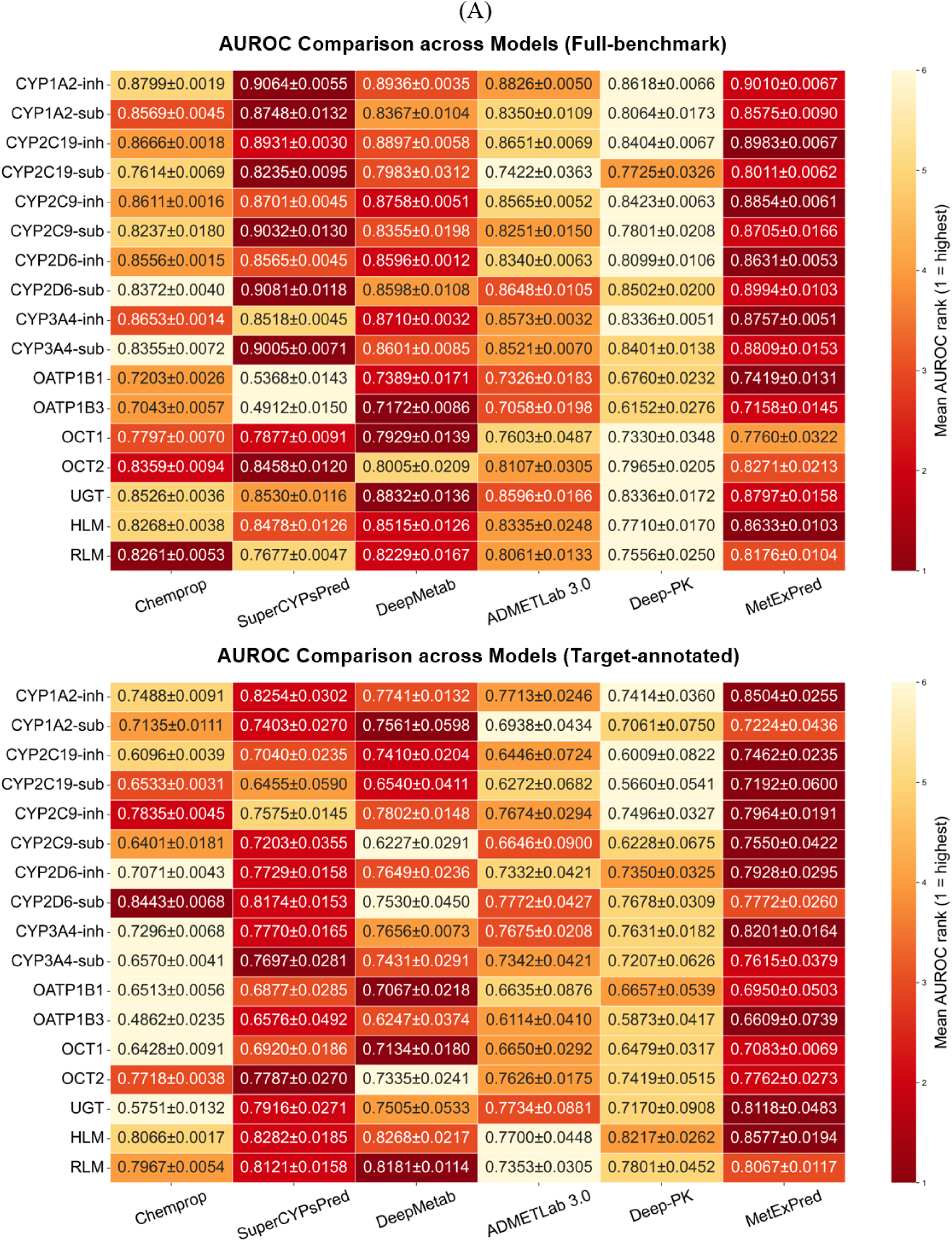
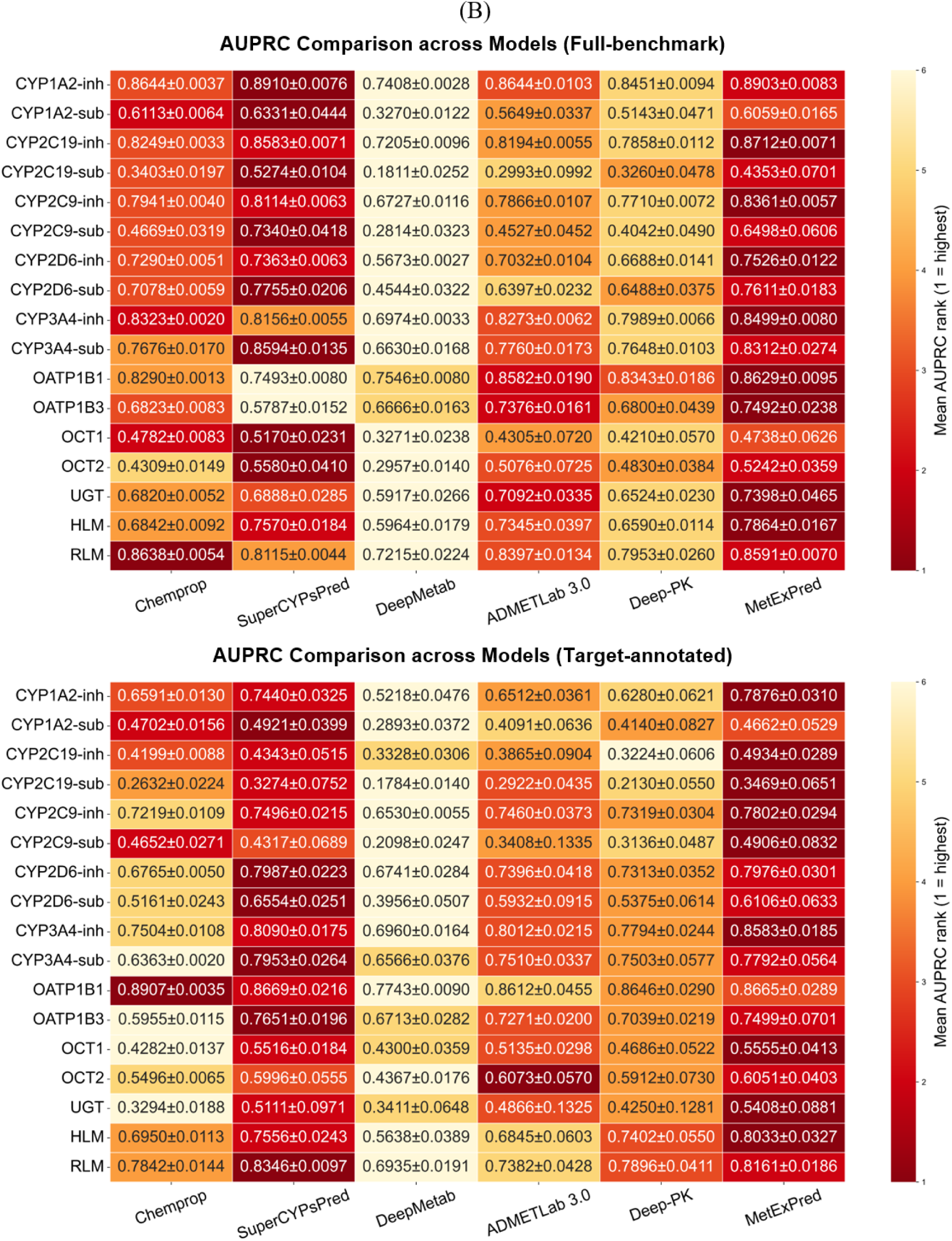
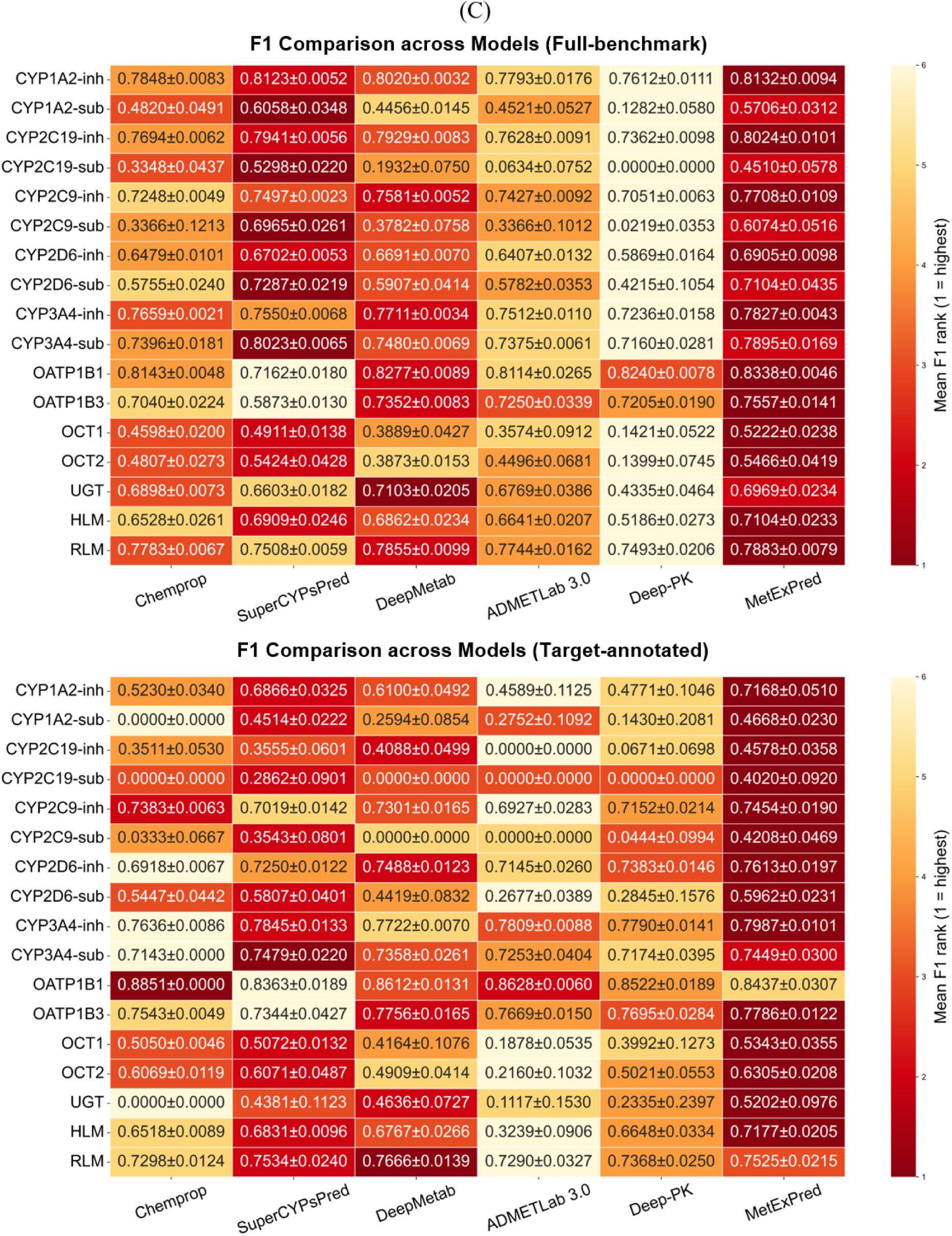
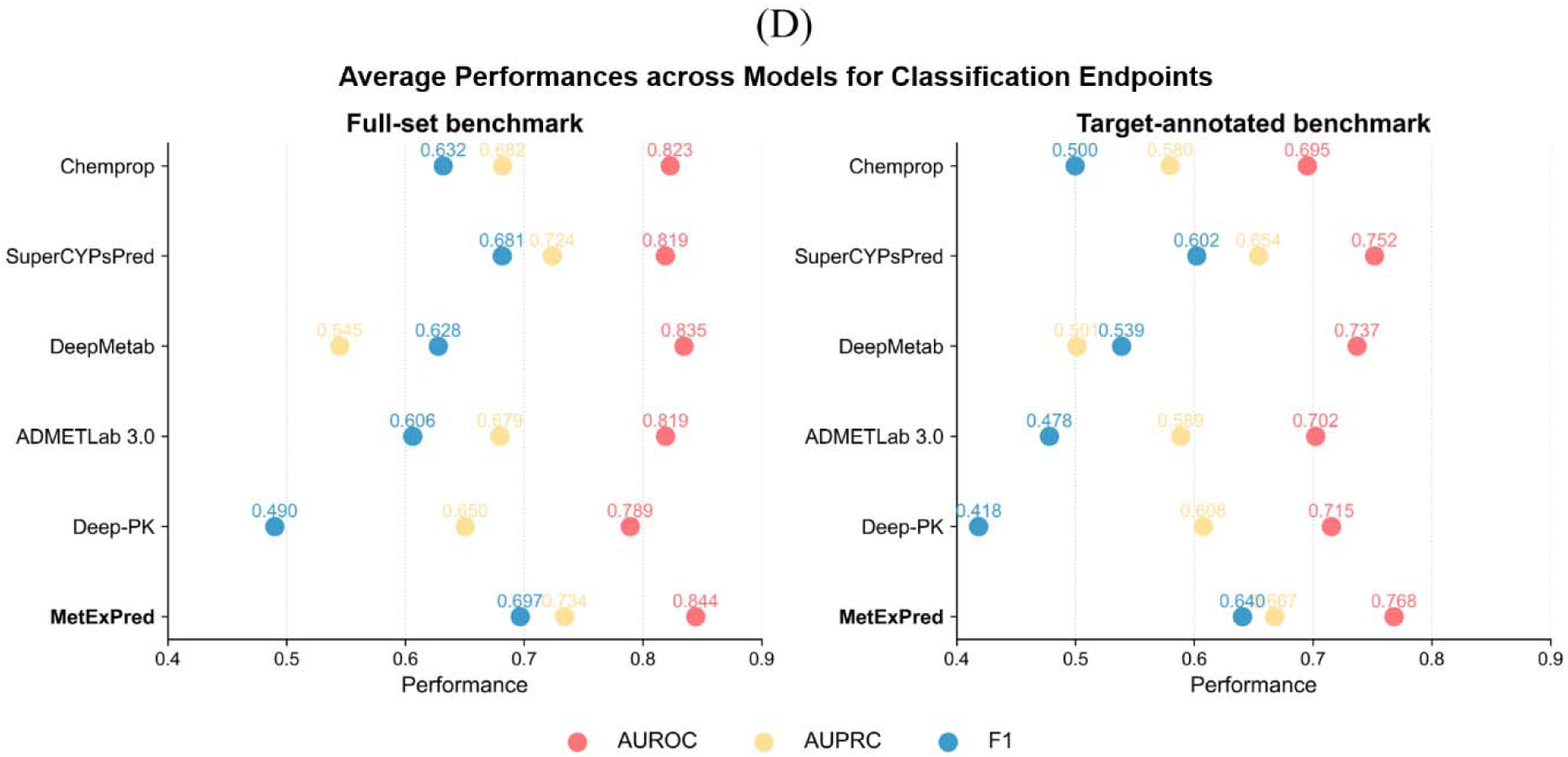
Performance comparison with existing baseline methods across classification endpoints. (A) Endpoint-level AUROC scores on the complete molecular benchmark and target-annotated subset. (B) Corresponding comparison of endpoint-level AUPRC score. (C) Corresponding comparison of endpoint-level F1 score. (D) Average performance across classification endpoints for two comparison groups. All methods were evaluated using the same curated data, split strategy, and evaluation protocol.

**Table 2.**
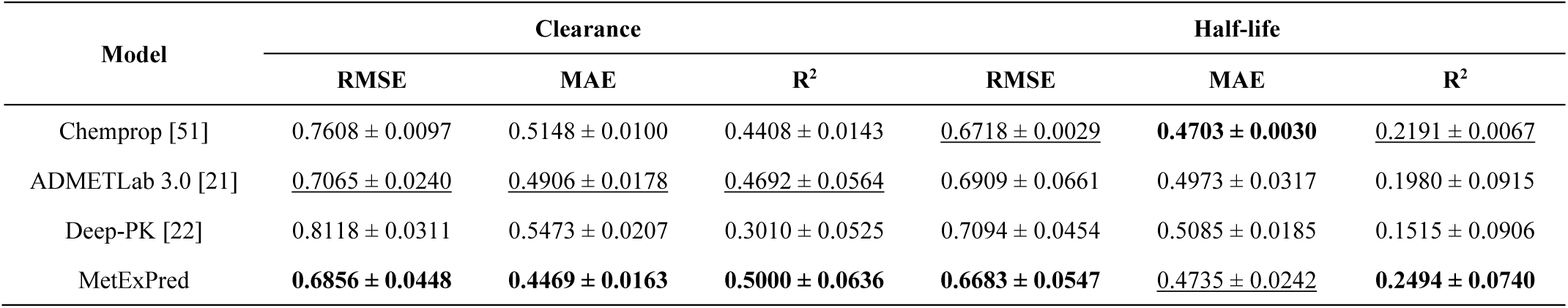
Performance comparison with baseline methods across excretion endpoints on the complete benchmark.

The same overall pattern was retained on the target-annotated subset. MetExPred remained competitive when only molecular representations were available and showed a modest additional benefit when protein representations were incorporated (Figure 4B and Table 3). The improvement was more evident for endpoints in which target-associated biological context contributed information beyond chemical structure. These results indicate that the proposed architecture does not depend on target availability to produce useful predictions, while still being able to exploit target information when it is present.

**Table 3.** Performance comparison with baseline methods across excretion endpoints on the target-annotated subset.

| Model | Clearance |  |  | Half-life |  |  |
| --- | --- | --- | --- | --- | --- | --- |
|  | RMSE | MAE | R<sup>2</sup> | RMSE | MAE | R<sup>2</sup> |
| Chemprop [51] | <u>0.5957 ± 0.0234</u> | <u>0.4333 ± 0.0059</u> | <u>0.4886 ± 0.0404</u> | 0.6518 ± 0.0074 | 0.4706 ± 0.0056 | 0.0483 ± 0.0215 |
| ADMETLab 3.0 [21] | 0.7139 ± 0.0691 | 0.5217 ± 0.0338 | 0.3354 ± 0.0480 | 0.6661 ± 0.0651 | 0.4788 ± 0.0297 | <u>0.1216 ± 0.0622</u> |
| Deep-PK [22] | 0.6339 ± 0.0446 | 0.4573 ± 0.0248 | 0.4687 ± 0.0698 | <u>0.6505 ± 0.0669</u> | <u>0.4705 ± 0.0317</u> | 0.1141 ± 0.1267 |
| MetExPred | **0.5883 ± 0.0233** | **0.4206 ± 0.0148** | **0.5438 ± 0.0291** | **0.6171 ± 0.0430** | **0.4480 ± 0.0180** | **0.2043 ± 0.0568** |

Note: For each evaluation metric, the best-performing value is highlighted in **bold**, and the second-best is indicated with underline for ease of comparison.

### 3.3 Ablation and model-design analysis

Detailed design analysis confirmed that the performance gains were attributable to complementary representations and adaptive fusion rather than to dimensionality alone. Compared with direct concatenation, linear attention, and convolutional neural network (CNN)-based attention, multi-view attention achieved the highest mean classification scores (AUROC 0.8444, AUPRC 0.7340, and F1 0.6966). It also produced the lowest errors for clearance (RMSE 0.6856 and MAE 0.4469) and half-life (RMSE 0.6683 and MAE 0.4735), while increasing the corresponding R-squared values to 0.5000 and 0.2494 (Figure 5A). These results indicate that jointly modeling view importance and cross-view dependencies is more effective than assigning fixed or locally derived contributions to heterogeneous embeddings.

**Figure 5.**
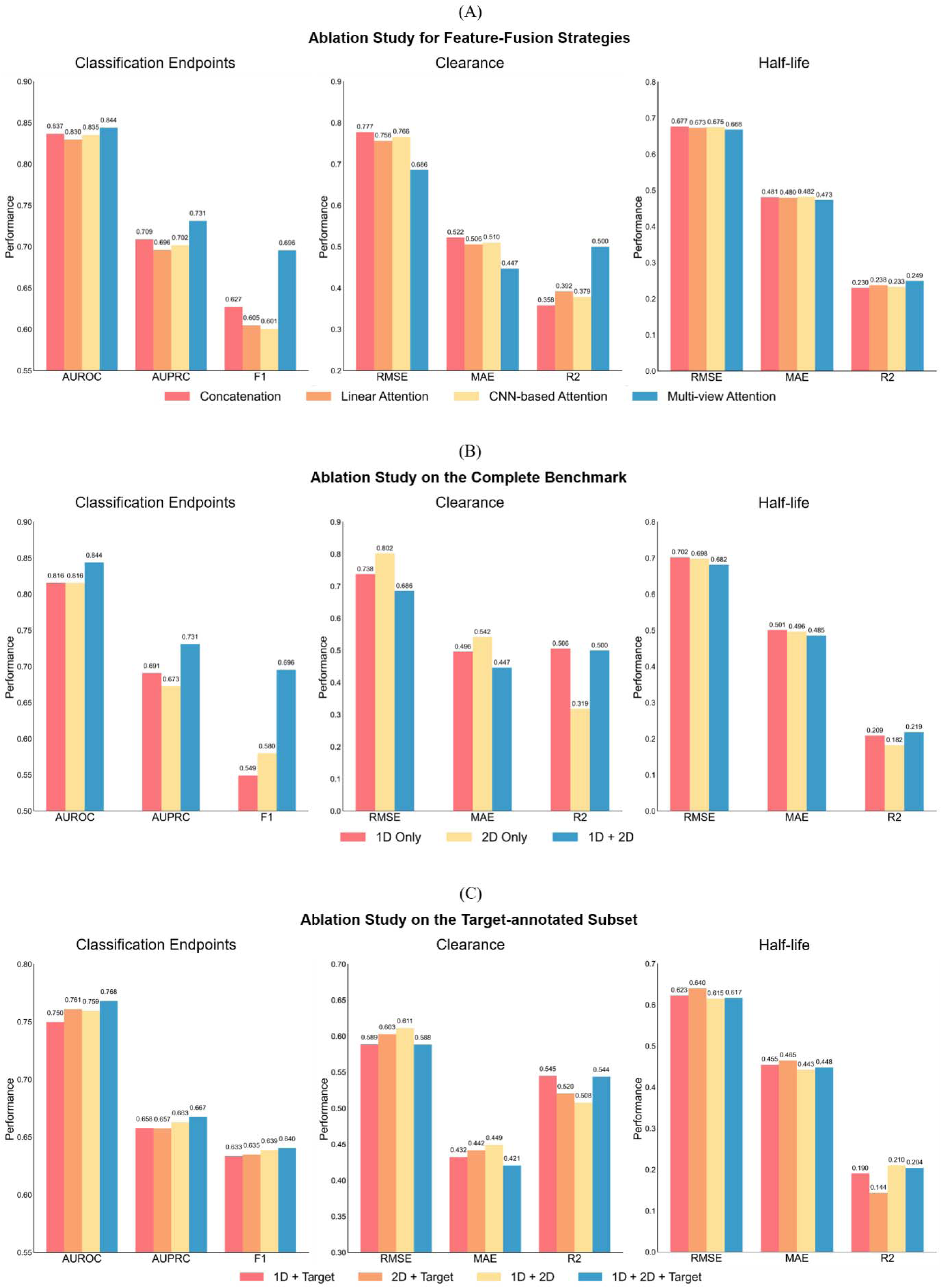
Ablation and model-design analysis. (A) Comparison of feature-fusion strategies for classification, clearance, and half-life prediction. (B) Contribution of 1D and 2D molecular representations on the complete benchmark. (C) Contribution of target information on the target-annotated subset.

The molecular-view ablation showed that 1D and 2D representations provided similar average AUROC values but differed more substantially in AUPRC, F1, and regression performance. ChemBERTa was particularly informative for continuous clearance and half-life outcomes, whereas D-MPNN contributed complementary local topology. Their integration improved all three mean classification metrics and reduced regression error relative to either molecular view alone (Figure 5B).

On the target-annotated subset, protein embeddings alone did not replace chemical representations. Instead, the complete 1D+2D+target configuration achieved the highest aggregate classification performance among the four target-only ablations, with mean AUROC, AUPRC, and F1 values of approximately 0.7681, 0.6675, and 0.6405, respectively. In regression endpoints, incorporating target information provided a modest improvement for clearance, whereas no consistent benefit was observed for half-life prediction (Figure 5C). Overall, these results suggest that protein representations provide complementary biological context rather than serving as a substitute for molecular structure, while their predictive contribution appears to be dependent on the specific endpoint.

### 3.4 DrugBank-based CYP profiling and literature-grounded case studies

The DrugBank-based case analysis first examined five compounds with diverse CYP inhibition profiles under molecular-only and target-aware prediction settings (Figure 6A–B). Because the five CYP inhibition models used endpoint-specific optimized decision thresholds, prediction probabilities were transformed using logit-based threshold normalization, with each endpoint-specific threshold mapped to a common value of 0.5. This transformation preserved the original positive/negative assignments and relative prediction ordering while enabling direct visualization across endpoints. The resulting values are therefore referred to as normalized prediction scores rather than calibrated probabilities.

**Figure 6.**
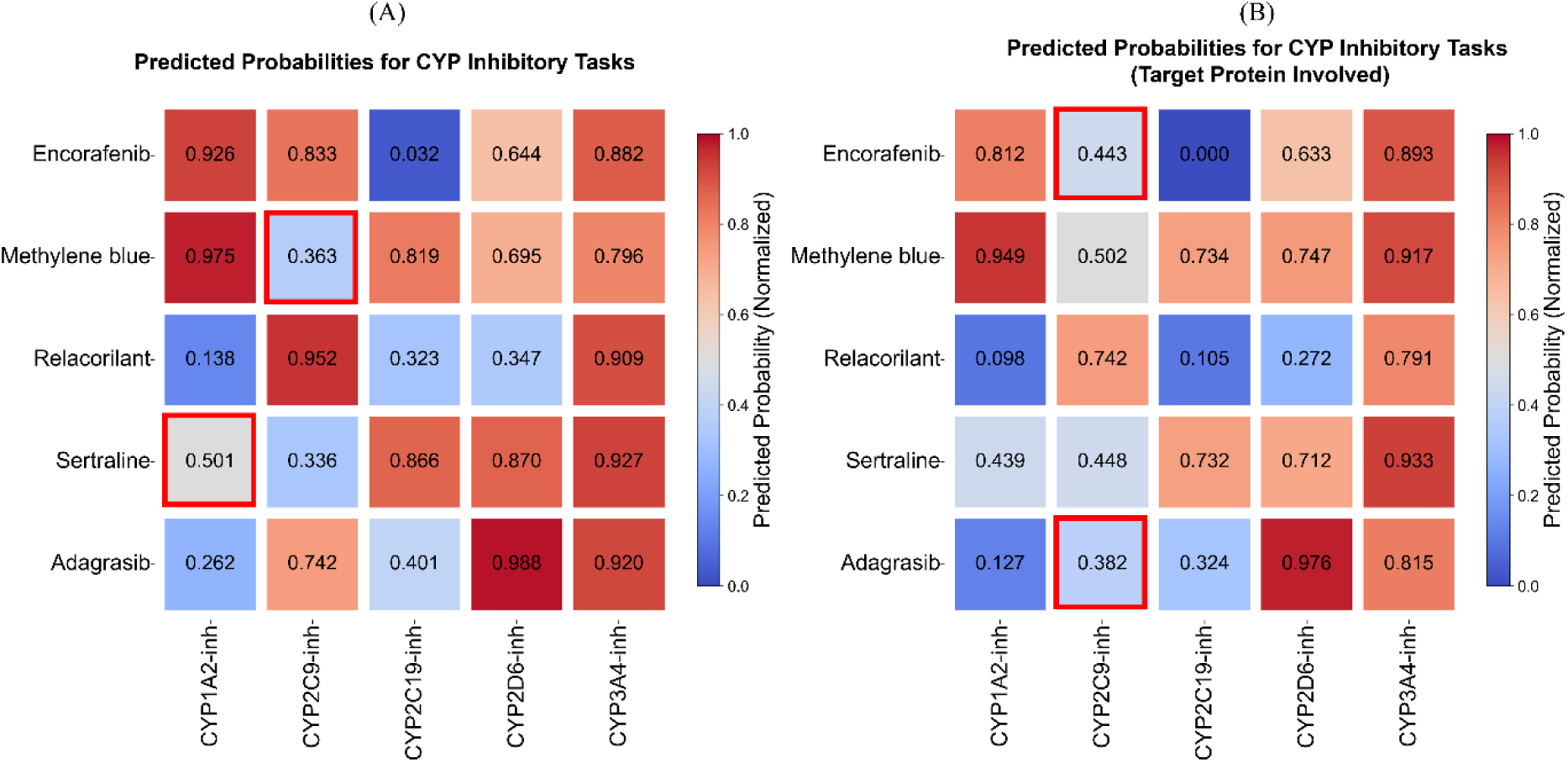
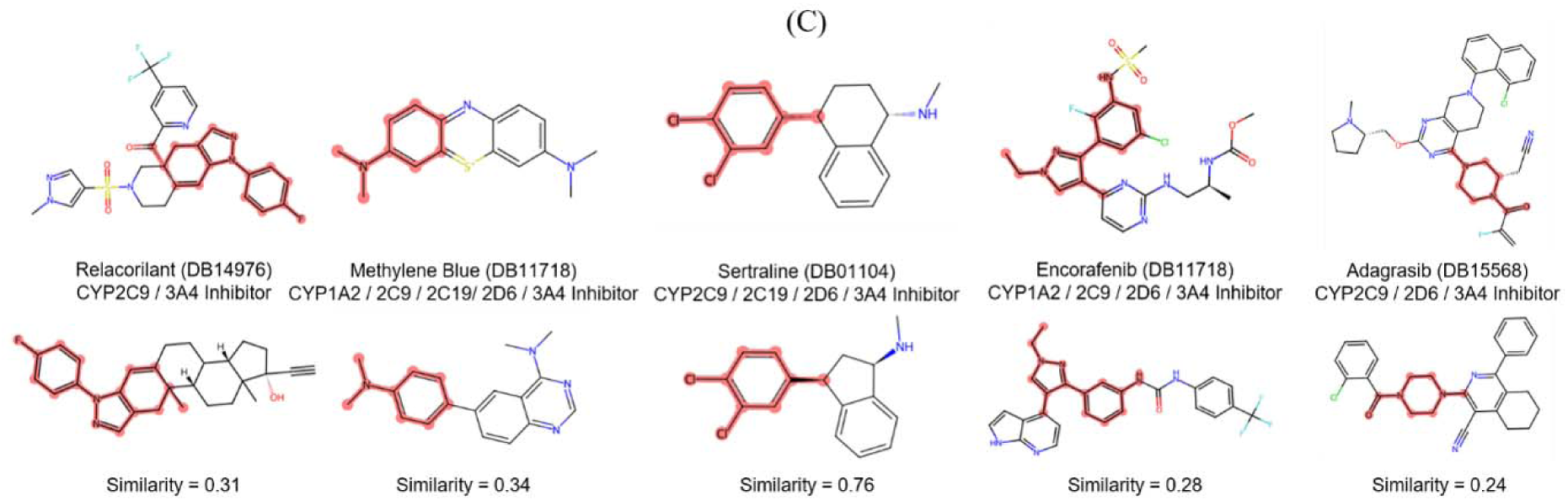
DrugBank-derived case studies of molecular-only and target-aware CYP inhibition prediction. (A) Normalized CYP inhibition prediction scores obtained using 1D and 2D molecular representations for five compounds selected from DrugBank. (B) Corresponding predictions after incorporating target-derived protein representations. Endpoint-specific decision thresholds were normalized to 0.5 for cross-endpoint visualization. Red boxes indicate representative discordant predictions relative to DrugBank annotations. (C) Maximum common substructure comparison between the selected compounds and their most similar training-set molecules, with Tanimoto similarity indicating structural familiarity.

Molecular-only predictions recovered most of the CYP inhibition patterns annotated in DrugBank for encorafenib, methylene blue, relacorilant, sertraline, and adagrasib. Incorporation of target-derived protein representations generally preserved the dominant prediction patterns while modifying several enzyme-specific scores. Consistent with the benchmark-level results, the effect of protein context was compound- and endpoint-dependent rather than uniformly beneficial. For example, target-aware modeling increased the normalized CYP2C9 inhibition score of methylene blue toward the positive side of the decision boundary, suggesting that biological context can supplement molecular information when the structural signal alone is less pronounced.

Several discordant predictions remained close to their endpoint-specific decision thresholds, including the molecular-only CYP1A2 prediction for sertraline and the target-aware CYP2C9 predictions for encorafenib and adagrasib. Maximum common substructure analysis further indicated that structural familiarity contributed to, but did not fully determine, prediction behavior (Figure 6C). Sertraline was predicted reliably in a relatively familiar chemical space, whereas relacorilant retained accurate predictions despite lower similarity to the training set. Conversely, methylene blue remained challenging despite moderate nearest-neighbor similarity. These observations indicate that structural similarity is better interpreted as a measure of chemical-space familiarity than as a direct surrogate for endpoint-specific prediction reliability. To extend the case analysis beyond the CYP-centered DrugBank examples, we further examined literature-grounded drug–endpoint pairs spanning a broader range of ME processes (Table 4). The model showed concordant predictions for most of the literature-supported classification cases, including CYP substrate and inhibitor activities, UGT substrate activity, and transporter-mediated disposition. Representative examples included the CYP1A2 substrate classification of fezolinetant [55], multi-enzyme substrate profiles of mavacamten [56] and deuruxolitinib, and OATP1B1/OATP1B3 inhibition by resmetirom[57]. At the same time, several discordant predictions were observed, particularly for selected transporter and CYP inhibition annotations of acoramidis [58] and arimoclomol [57, 59], indicating that individual endpoint predictions should still be interpreted together with confidence and endpoint-specific evidence.

**Table 4.**
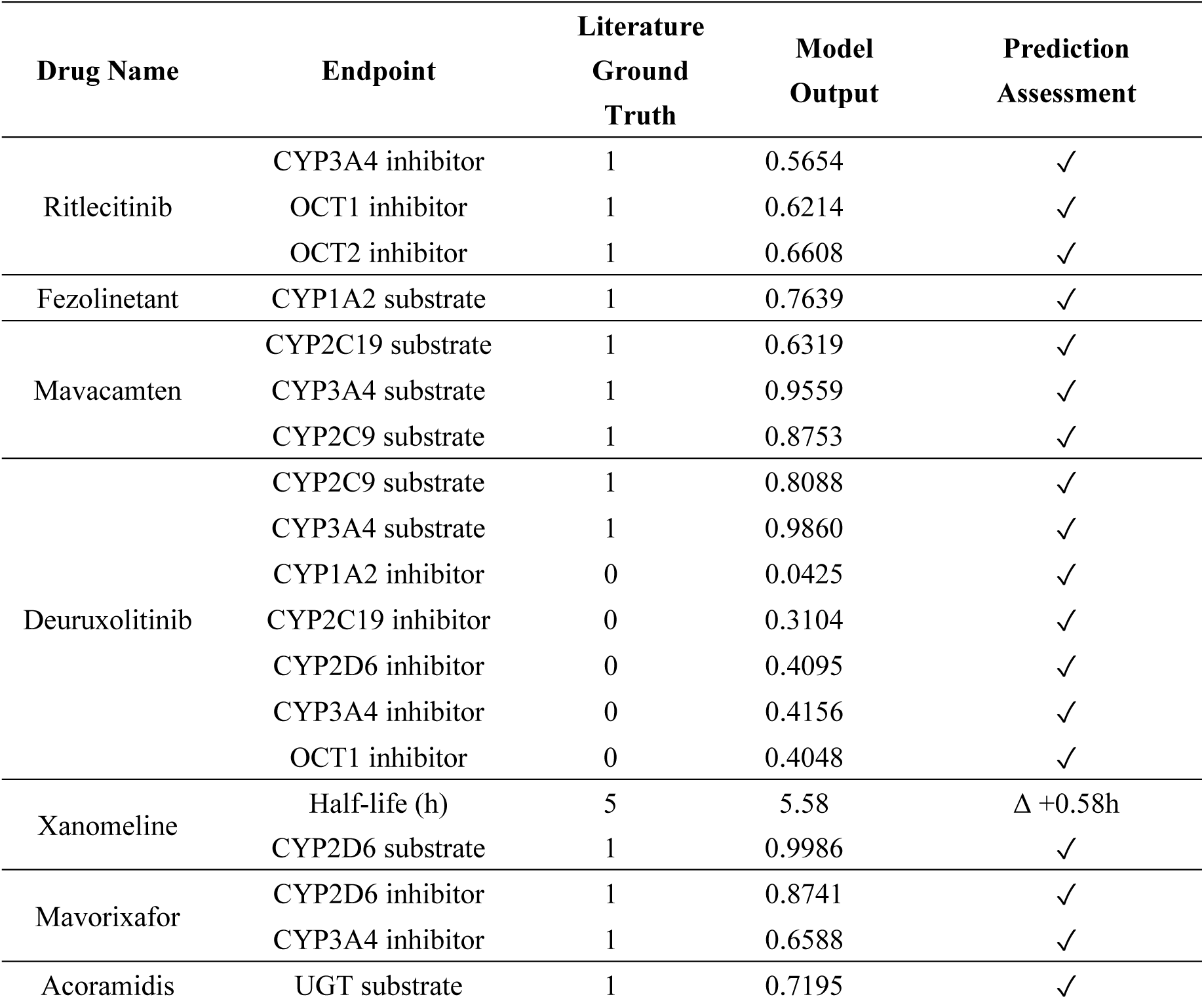
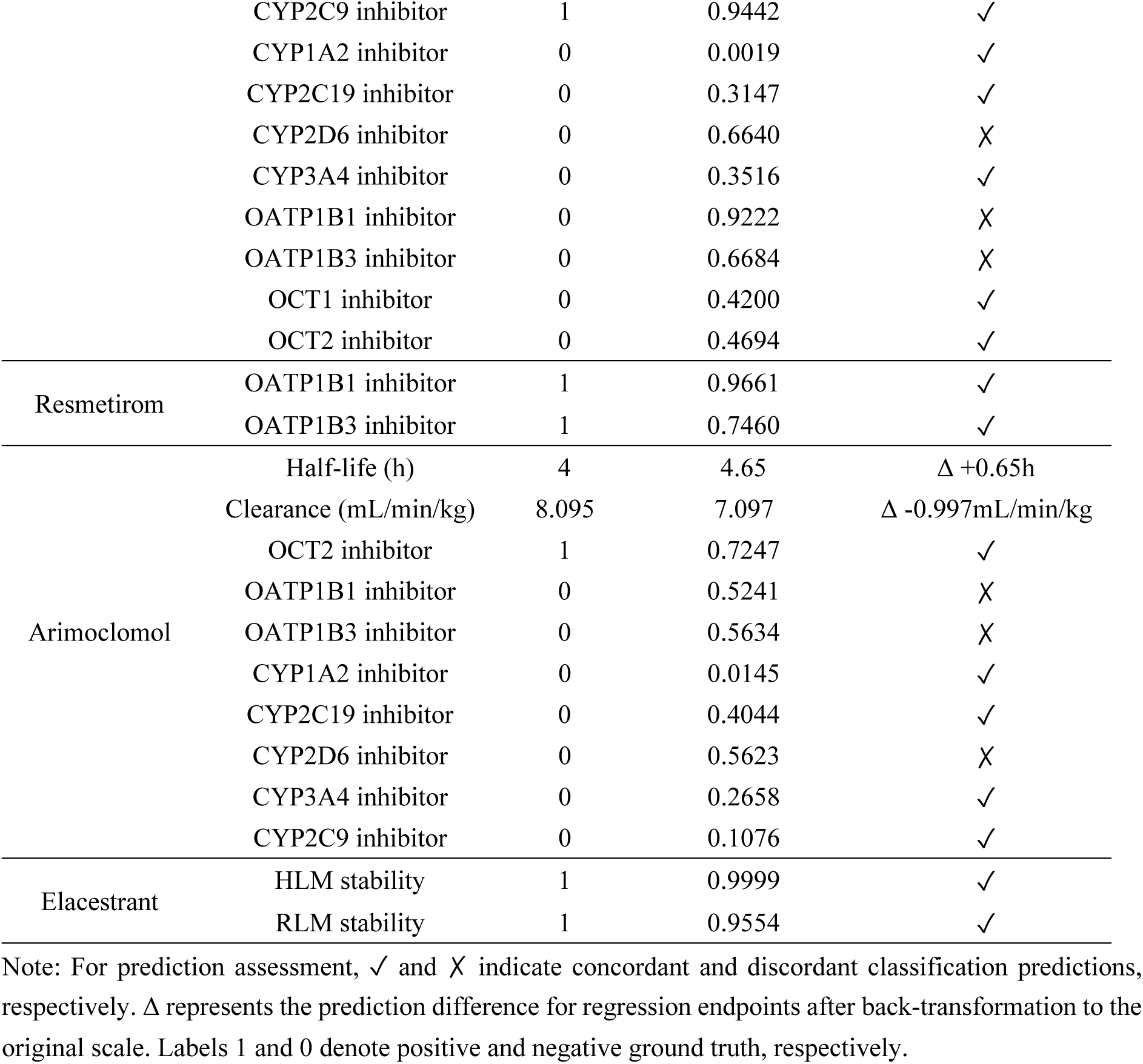
Literature-grounded case studies of MetExPred across representative metabolism and excretion endpoints.

The literature-grounded regression cases provided complementary quantitative examples. For xanomeline, the predicted half-life of approximately 5.58 h was close to the reported value of 5.0 h [58], while the predicted half-life of arimoclomol was approximately 4.65 h compared with a reported value of 4.0 h [59]. For arimoclomol clearance, the predicted value of 7.10 mL/min/kg was also close to the literature-derived value of 8.10 mL/min/kg. [57]Together, the DrugBank-based visualization and literature-grounded cases illustrate two complementary aspects of MetExPred: the former provides a controlled comparison of molecular-only and target-aware prediction behavior, whereas the latter demonstrates the applicability of the framework across heterogeneous metabolism and excretion endpoints.

## 4 Discussion

This study addresses ME prediction at both data and representation levels. The curated benchmark not only expands shared CYP and elimination-related pharmacokinetic tasks, but also adds underrepresented UGT and transporter endpoints, harmonizing them within one evaluation framework. MetExPred then combines molecular language, molecular graph and target-derived protein representations through masked multi-view attention. The overall benchmark, baseline comparison and ablation results consistently indicate that these views are complementary. In particular, the model retains strong performance without target annotations and obtains additional benefits from protein context on the target-annotated subset.

The results indicate that molecular sequence and graph representations constitute the core predictive basis of MetExPred, whereas target annotations provide optional biological context when available. This design is consistent with the intended use of the framework in both molecular-only and target-enhanced settings. The ChEMBL-derived targets include diverse pharmacological proteins and therefore should not be interpreted as direct mechanistic evidence that a particular protein mediates the corresponding ME endpoint [48]. Instead, these annotations summarize experimentally reported biological interaction profiles and polypharmacological properties that provide information complementary to molecular structure. Although the gains from target-derived protein representations were modest and varied across endpoints (Figure 5C), intended targets and experimentally measured compound-target activities may already be available during hit identification and lead optimization [38, 39]. Including these annotations therefore allows the benchmark to more fully reflect the range of information that may be available during early drug development and to evaluate when target context adds value beyond molecular structure, without assuming a uniform benefit across endpoints. The protein-aware masking strategy enables the same architecture to process compounds with or without target annotations by excluding missing protein representations from attention-based fusion, thereby preventing placeholder vectors from contributing to the final prediction.

The differences observed among classification and regression tasks further highlight the heterogeneous nature of ME prediction. CYP and transporter classification endpoints are often associated with recurring structural motifs and physicochemical patterns that can be learned from molecular representations. In contrast, clearance and half-life are integrated pharmacokinetic outcomes influenced by multiple molecular, biological and physiological factors. The comparatively lower R-squared value for half-life may therefore reflect its dependence on distribution, plasma protein binding, metabolism, excretion and variability in experimental conditions [4, 6, 7, 12]. Although a unified architecture provides a consistent framework for modeling diverse ME properties, endpoint-specific evaluation and interpretation remain necessary. Several limitations should also be acknowledged. The benchmark integrates data from heterogeneous public and curated sources, and residual differences in assay conditions, activity thresholds and annotation quality may remain after harmonization [21–23, 48]. In addition, the current use of random splitting may yield optimistic estimates for structurally novel compounds, and future evaluation using scaffold-based, temporal and independent external test sets will be important [60]. Target annotations are incomplete and biased toward well-characterized compounds and proteins, while mean pooling may obscure distinct contributions from individual targets. The current framework also does not explicitly model direct compound-protein interactions or causal relationships among metabolic activity, transporter function, clearance and half-life. Finally, the present case studies provide illustrative examples rather than a comprehensive evaluation of prediction uncertainty and applicability domains.

MetExPred is therefore intended as an early-stage prioritization framework rather than a replacement for experimental ADMET assays or clinical pharmacokinetic assessment [1–3]. It can support the identification of CYP and transporter liabilities, prioritization of compounds for microsomal stability experiments and preliminary estimation of clearance and half-life. Its ability to operate without target annotations also makes it applicable to novel compounds for which biological information is unavailable, while the curated target data provide an additional resource for investigating whether biological context improves individual predictions. Future work should incorporate uncertainty calibration, applicability-domain assessment, scaffold-based and external validation, more explicit compound-protein interaction modeling and cross-endpoint learning. A user-facing platform could further provide endpoint-level predictions, confidence estimates, target availability, attention weights and structurally similar reference compounds to facilitate transparent interpretation and practical application.

## 5 Conclusion

This study established a comprehensive benchmark covering 17 metabolism-related classification endpoints and two excretion-related regression endpoints and developed MetExPred for prediction across these tasks. MetExPred showed the strongest average performance among the evaluated baselines and remained applicable in both molecular-only and target-aware settings. These findings support its use for early-stage assessment of drug metabolism and excretion properties.

## Data and Code Availability

Benchmark datasets of this study are available in the following sources: Deep-PK at https://biosig.lab.uq.edu.au/deeppk/data, reference number [22]; admetSAR 3.0 at https://lmmd.ecust.edu.cn/admetsar3/resource/ADME.php, reference number [23]; Therapeutic Data Commons at https://tdcommons.ai/single_pred_tasks/adme, reference number [40]; CMDM at https://doi.org/10.1038/s41597-025-05753-8, reference number [41]; ADME@NCATS at https://opendata.ncats.nih.gov/adme/data, reference number [42]; PharmaBench at https://github.com/mindrank-ai/PharmaBench, reference number [43]; IsTransbase at https://istransbase.scbdd.com/database, reference number [44]; Maom Lab at https://doi.org/10.1021/acs.chemrestox.2c00207, reference number [45]; DDPD at http://www.inbirg.com/ddpd/download, reference number [46]; DrugBank at https://go.drugbank.com, reference number [48]; and ChEMBL 36 at https://www.ebi.ac.uk/chembl, reference number [49]. Source code and pretrained checkpoints used in this study are available at the GitHub repository: https://github.com/angel2002hyx-lgtm/MetExPred.

## Author Contributions

Y.H. conceived the study, curated data, performed formal analyses, generated visualizations, and wrote the original draft. J.W. conceived the study, developed methodology, performed formal analyses, and wrote the original draft. G.W. contributed to data curation, wrote parts of the original draft, and participated in review and editing. D.D. assisted with the methodology of case studies and revised the manuscript. Y-C-D.L. reviewed and edited the manuscript. H-Y.H. contributed to methodology development, supervised the work, and critically revised the manuscript. H-D.H. provided overall supervision, led conceptualization, offered methodological guidance, and approved the final revision. All authors read and approved the final manuscript.

## Acknowledgements

This research benefited significantly from the interdisciplinary research environment and cut-ting-edge instrumentation maintained by the Computational Platform of Warshel Institute for Computational Biology. Their ongoing commitment to research infrastructure development has been crucial to our scientific endeavors. Authors are also grateful to the library of The Chinese University of Hong Kong, Shenzhen for providing effective database service.

