## Supplementary Files for "MetExPred: A Comprehensive Prediction Framework with Protein-Context-Aware Multi-view Learning for Drug Metabolism and Excretion"

Supplementary File

**Table S1.** Train, validation and test splits for MetExPred Benchmark Dataset

| Endpoint | Training Set | Validation Set | Test Set |
|---|---|---|---|
| CYP1A2 Inhibitor | 19,481 | 2,435 | 2,436 |
| CYP1A2 Substrate | 3,069 | 384 | 384 |
| CYP2C19 Inhibitor | 19,444 | 2,430 | 2,431 |
| CYP2C19 Substrate | 3,012 | 377 | 377 |
| CYP2C9 Inhibitor | 22,861 | 2,858 | 2,858 |
| CYP2C9 Substrate | 3,446 | 431 | 431 |
| CYP2D6 Inhibitor | 23,741 | 2,968 | 2,968 |
| CYP2D6 Substrate | 3,458 | 432 | 433 |
| CYP3A4 Inhibitor | 26,313 | 3,289 | 3,290 |
| CYP3A4 Substrate | 4,047 | 506 | 506 |
| HLM Stability | 4,818 | 602 | 603 |
| RLM Stability | 4,769 | 596 | 597 |
| UGTs Substrate | 2,131 | 266 | 267 |
| OATP1B1 Inhibitor | 3,260 | 408 | 408 |
| OATP1B3 Inhibitor | 3,190 | 399 | 399 |
| OCT1 Inhibitor | 2,702 | 338 | 338 |
| OCT2 Inhibitor | 3,100 | 388 | 388 |
| Clearance | 4,616 | 577 | 577 |
| Half-life | 2,096 | 262 | 263 |

**Table S2.** Target Annotation Information for MetExPred Benchmark Dataset

| Endpoint | Targets | Molecules with Target(s) | Molecule-Target Pairs |
|---|---|---|---|
| CYP1A2 Inhibitor | 830 | 1,351 | 6,390 |
| CYP1A2 Substrate | 754 | 860 | 5,153 |
| CYP2C19 Inhibitor | 839 | 1,367 | 6,409 |
| CYP2C19 Substrate | 756 | 853 | 5,174 |
| CYP2C9 Inhibitor | 1,055 | 2,890 | 11,438 |
| CYP2C9 Substrate | 751 | 902 | 5,275 |
| CYP2D6 Inhibitor | 1,051 | 3,009 | 11,717 |
| CYP2D6 Substrate | 752 | 908 | 5,273 |
| CYP3A4 Inhibitor | 1,074 | 3,527 | 12,274 |
| CYP3A4 Substrate | 785 | 984 | 5,612 |
| HLM Stability | 483 | 2,067 | 3,660 |
| RLM Stability | 1,129 | 1,759 | 4,872 |
| UGTs Substrate | 776 | 847 | 5,343 |
| OATP1B1 Inhibitor | 818 | 965 | 5,232 |
| OATP1B3 Inhibitor | 817 | 963 | 5,212 |
| OCT1 Inhibitor | 814 | 936 | 5,102 |
| OCT2 Inhibitor | 826 | 1,045 | 5,485 |
| Clearance | 1,004 | 2,471 | 7,004 |
| Half-life | 758 | 923 | 4,990 |

**Table S3.** AUROC scores for 1D+2D embedding combinations across classification endpoints

| Endpoint | 1D TF-IDF | 1D ChemBERTa | 1D MoLFormer | 2D Morgan FP | 2D D-MPNN | TF-IDF + Morgan FP | ChemBERTa + Morgan FP |
|---|---|---|---|---|---|---|---|
| CYP1A2 Inhibitor | 0.8592 ± 0.0068 | 0.8892 ± 0.0077 | 0.8606 ± 0.0086 | 0.8700 ± 0.0107 | **0.8947 ± 0.0055** | 0.8656 ± 0.0048 | 0.8715 ± 0.0045 |
| CYP1A2 Substrate | 0.7817 ± 0.0319 | 0.8283 ± 0.0214 | 0.8089 ± 0.0230 | 0.8326 ± 0.0258 | 0.8106 ± 0.0337 | <u>0.8434 ± 0.0124</u> | **0.8559 ± 0.0064** |
| CYP2C19 Inhibitor | 0.8526 ± 0.0058 | 0.8803 ± 0.0076 | 0.8568 ± 0.0039 | 0.8662 ± 0.0077 | 0.8818 ± 0.0089 | 0.8621 ± 0.0073 | 0.8714 ± 0.0102 |
| CYP2C19 Substrate | 0.7756 ± 0.0317 | 0.8038 ± 0.0216 | 0.7767 ± 0.0408 | **0.8300 ± 0.0280** | 0.7476 ± 0.0507 | <u>0.8056 ± 0.0313</u> | 0.8139 ± 0.0268 |
| CYP2C9 Inhibitor | 0.8429 ± 0.0064 | 0.8672 ± 0.0030 | 0.8448 ± 0.0053 | 0.8483 ± 0.0037 | 0.8736 ± 0.0051 | 0.8482 ± 0.0042 | 0.8575 ± 0.0054 |
| CYP2C9 Substrate | 0.8183 ± 0.0159 | 0.8197 ± 0.0147 | 0.7810 ± 0.0573 | 0.8623 ± 0.0219 | 0.7879 ± 0.0209 | <u>0.8701 ± 0.0278</u> | **0.8736 ± 0.0275** |
| CYP2D6 Inhibitor | 0.8161 ± 0.0087 | 0.8458 ± 0.0047 | 0.8135 ± 0.0049 | 0.8168 ± 0.0107 | <u>0.8635 ± 0.0080</u> | 0.8232 ± 0.0080 | 0.8327 ± 0.0067 |
| CYP2D6 Substrate | 0.8436 ± 0.0025 | 0.8526 ± 0.0151 | 0.8533 ± 0.0130 | 0.8733 ± 0.0152 | 0.8308 ± 0.0193 | 0.8821 ± 0.0154 | <u>0.8821 ± 0.0145</u> |
| CYP3A4 Inhibitor | 0.8281 ± 0.0068 | 0.8550 ± 0.0035 | 0.8328 ± 0.0043 | 0.8424 ± 0.0056 | **0.8757 ± 0.0066** | 0.8425 ± 0.0062 | 0.8511 ± 0.0068 |
| CYP3A4 Substrate | 0.8611 ± 0.0133 | 0.8697 ± 0.0230 | 0.8458 ± 0.0182 | **0.8861 ± 0.0158** | 0.8524 ± 0.0107 | <u>0.8761 ± 0.0085</u> | 0.8721 ± 0.0088 |
| OATP1B1 Inhibitor | 0.6666 ± 0.0433 | 0.6584 ± 0.0332 | 0.6954 ± 0.0283 | 0.6176 ± 0.0205 | 0.6880 ± 0.0105 | 0.6609 ± 0.0235 | 0.6565 ± 0.0228 |
| OATP1B3 Inhibitor | 0.6762 ± 0.0253 | 0.6008 ± 0.0243 | 0.6745 ± 0.0187 | 0.5612 ± 0.0187 | **0.7232 ± 0.0242** | 0.5952 ± 0.0288 | 0.5766 ± 0.0377 |
| OCT1 Inhibitor | 0.7350 ± 0.0542 | 0.7745 ± 0.0250 | 0.7768 ± 0.0403 | 0.7482 ± 0.0289 | 0.7737 ± 0.0276 | 0.7729 ± 0.0250 | <u>0.7776 ± 0.0254</u> |
| OCT2 Inhibitor | 0.7656 ± 0.0405 | 0.8150 ± 0.0363 | 0.7855 ± 0.0199 | 0.8100 ± 0.0290 | 0.8192 ± 0.0418 | **0.8197 ± 0.0070** | 0.8182 ± 0.0091 |
| UGTs Substrate | 0.7971 ± 0.0385 | <u>0.8592 ± 0.0263</u> | 0.8125 ± 0.0251 | 0.8023 ± 0.0227 | 0.8224 ± 0.0285 | 0.8243 ± 0.0310 | 0.8562 ± 0.0251 |
| HLM Stability | 0.8008 ± 0.0307 | <u>0.8523 ± 0.0157</u> | 0.8100 ± 0.0159 | 0.8357 ± 0.0131 | 0.8080 ± 0.0143 | 0.8376 ± 0.0156 | 0.8398 ± 0.0183 |
| RLM Stability | 0.7894 ± 0.0133 | 0.7976 ± 0.0173 | 0.7517 ± 0.0213 | 0.7933 ± 0.0045 | <u>0.8142 ± 0.0161</u> | 0.7879 ± 0.0203 | 0.7924 ± 0.0180 |

Note: For each evaluation metric, the best-performing value is highlighted in **bold**, and the second-best is indicated with <u>underline</u> for ease of comparison.

**Table S3** (continued)

| Endpoint | MoLFormer + Morgan FP | TF-IDF + D-MPNN | ChemBERTa + D-MPNN | MoLFormer + D-MPNN |
|---|---|---|---|---|
| CYP1A2 Inhibitor | 0.8716 ± 0.0023 | 0.8849 ± 0.0059 | <u>0.8939 ± 0.0054</u> | 0.8894 ± 0.0056 |
| CYP1A2 Substrate | 0.8481 ± 0.0066 | 0.8287 ± 0.0144 | 0.8414 ± 0.0195 | 0.8420 ± 0.0290 |
| CYP2C19 Inhibitor | 0.8659 ± 0.0083 | 0.8792 ± 0.0084 | **0.8923 ± 0.0048** | <u>0.8857 ± 0.0045</u> |
| CYP2C19 Substrate | 0.8050 ± 0.0205 | 0.7472 ± 0.0540 | 0.7832 ± 0.0495 | 0.7642 ± 0.0391 |
| CYP2C9 Inhibitor | 0.8525 ± 0.0066 | <u>0.8755 ± 0.0080</u> | **0.8823 ± 0.0038** | 0.8750 ± 0.0081 |
| CYP2C9 Substrate | 0.8673 ± 0.0270 | 0.8384 ± 0.0223 | 0.8619 ± 0.0141 | 0.8455 ± 0.0176 |
| CYP2D6 Inhibitor | 0.8286 ± 0.0049 | 0.8554 ± 0.0055 | **0.8635 ± 0.0023** | 0.8592 ± 0.0033 |
| CYP2D6 Substrate | 0.8800 ± 0.0135 | **0.8822 ± 0.0125** | 0.8736 ± 0.0055 | 0.8677 ± 0.0118 |
| CYP3A4 Inhibitor | 0.8468 ± 0.0060 | 0.8696 ± 0.0070 | <u>0.8730 ± 0.0049</u> | 0.8715 ± 0.0040 |
| CYP3A4 Substrate | 0.8632 ± 0.0085 | 0.8565 ± 0.0112 | 0.8538 ± 0.0103 | 0.8570 ± 0.0186 |
| OATP1B1 Inhibitor | 0.6739 ± 0.0182 | **0.7336 ± 0.0166** | <u>0.7325 ± 0.0155</u> | 0.7150 ± 0.0081 |
| OATP1B3 Inhibitor | 0.6844 ± 0.0245 | 0.7046 ± 0.0175 | <u>0.7168 ± 0.0133</u> | 0.7161 ± 0.0209 |
| OCT1 Inhibitor | **0.7807 ± 0.0275** | 0.7619 ± 0.0477 | 0.7775 ± 0.0370 | 0.7723 ± 0.0455 |
| OCT2 Inhibitor | 0.8135 ± 0.0179 | 0.8107 ± 0.0300 | 0.8165 ± 0.0191 | <u>0.8192 ± 0.0246</u> |
| UGTs Substrate | 0.8433 ± 0.0264 | 0.8493 ± 0.0290 | **0.8873 ± 0.0143** | 0.8557 ± 0.0271 |
| HLM Stability | 0.8407 ± 0.0158 | 0.8287 ± 0.0267 | **0.8574 ± 0.0156** | 0.8220 ± 0.0225 |
| RLM Stability | 0.7834 ± 0.0217 | 0.8023 ± 0.0063 | **0.8224 ± 0.0096** | 0.8029 ± 0.0169 |

Note: For each evaluation metric, the best-performing value is highlighted in **bold**, and the second-best is indicated with <u>underline</u> for ease of comparison.

**Table S4.** AUPRC scores for 1D+2D embedding combinations across classification endpoints

| Endpoint | 1D TF-IDF | 1D ChemBERTa | 1D MoLFormer | 2D Morgan FP | 2D D-MPNN | TF-IDF + Morgan FP | ChemBERTa + Morgan FP |
|---|---|---|---|---|---|---|---|
| CYP1A2 Inhibitor | 0.8373 ± 0.0089 | 0.8704 ± 0.0041 | 0.8427 ± 0.0084 | 0.8426 ± 0.0129 | <u>0.8829 ± 0.0085</u> | 0.8408 ± 0.0039 | 0.8558 ± 0.0041 |
| CYP1A2 Substrate | 0.4608 ± 0.0591 | 0.5172 ± 0.0674 | 0.4919 ± 0.0300 | 0.5771 ± 0.0603 | 0.4644 ± 0.0766 | 0.5992 ± 0.0599 | **0.6242 ± 0.0514** |
| CYP2C19 Inhibitor | 0.8081 ± 0.0169 | 0.8430 ± 0.0120 | 0.8156 ± 0.0133 | 0.8217 ± 0.0125 | 0.8442 ± 0.0175 | 0.8176 ± 0.0100 | 0.8305 ± 0.0158 |
| CYP2C19 Substrate | 0.4129 ± 0.0625 | 0.3844 ± 0.0903 | 0.3543 ± 0.0564 | 0.5251 ± 0.0680 | 0.2932 ± 0.0480 | 0.4873 ± 0.0442 | **0.4964 ± 0.0706** |
| CYP2C9 Inhibitor | 0.7694 ± 0.0100 | 0.8057 ± 0.0072 | 0.7769 ± 0.0143 | 0.7755 ± 0.0096 | 0.8137 ± 0.0098 | 0.7778 ± 0.0098 | 0.7914 ± 0.0077 |
| CYP2C9 Substrate | 0.4826 ± 0.0346 | 0.5141 ± 0.0206 | 0.4652 ± 0.1228 | **0.6201 ± 0.0580** | 0.4119 ± 0.0687 | 0.6099 ± 0.0410 | 0.6056 ± 0.0542 |
| CYP2D6 Inhibitor | 0.6743 ± 0.0211 | 0.7287 ± 0.0067 | 0.6773 ± 0.0198 | 0.6870 ± 0.0265 | **0.7552 ± 0.0161** | 0.6882 ± 0.0125 | 0.7046 ± 0.0133 |
| CYP2D6 Substrate | 0.6302 ± 0.0336 | 0.6520 ± 0.0763 | 0.6578 ± 0.0444 | 0.6910 ± 0.0422 | 0.5915 ± 0.0320 | <u>0.7103 ± 0.0442</u> | **0.7211 ± 0.0254** |
| CYP3A4 Inhibitor | 0.7960 ± 0.0104 | 0.8287 ± 0.0061 | 0.7991 ± 0.0052 | 0.8192 ± 0.0075 | **0.8487 ± 0.0059** | 0.8122 ± 0.0084 | 0.8194 ± 0.0100 |
| CYP3A4 Substrate | 0.8089 ± 0.0134 | 0.8222 ± 0.0468 | 0.7825 ± 0.0343 | **0.8364 ± 0.0375** | 0.7864 ± 0.0313 | <u>0.8244 ± 0.0139</u> | 0.8158 ± 0.0154 |
| OATP1B1 Inhibitor | 0.8025 ± 0.0399 | 0.8130 ± 0.0358 | 0.8145 ± 0.0440 | 0.7882 ± 0.0329 | 0.8206 ± 0.0202 | 0.8196 ± 0.0094 | 0.8182 ± 0.0139 |
| OATP1B3 Inhibitor | 0.7116 ± 0.0394 | 0.6724 ± 0.0400 | 0.7230 ± 0.0164 | 0.6280 ± 0.0439 | **0.7540 ± 0.0328** | 0.6602 ± 0.0317 | 0.6470 ± 0.0449 |
| OCT1 Inhibitor | 0.4371 ± 0.0951 | **0.4679 ± 0.0547** | 0.4493 ± 0.0823 | 0.4487 ± 0.0835 | <u>0.4629 ± 0.0664</u> | 0.4435 ± 0.0302 | 0.4455 ± 0.0366 |
| OCT2 Inhibitor | 0.4708 ± 0.0725 | <u>0.5188 ± 0.1048</u> | 0.4735 ± 0.0360 | 0.5067 ± 0.0678 | **0.5459 ± 0.0873** | 0.4763 ± 0.0111 | 0.4776 ± 0.0139 |
| UGTs Substrate | 0.6103 ± 0.0767 | <u>0.7071 ± 0.0610</u> | 0.6340 ± 0.0250 | 0.6419 ± 0.0315 | 0.6258 ± 0.0573 | 0.6489 ± 0.0328 | 0.6727 ± 0.0264 |
| HLM Stability | 0.6927 ± 0.0373 | <u>0.7633 ± 0.0181</u> | 0.6952 ± 0.0164 | 0.7328 ± 0.0148 | 0.6791 ± 0.0520 | 0.7432 ± 0.0207 | 0.7452 ± 0.0251 |
| RLM Stability | 0.8211 ± 0.0179 | 0.8374 ± 0.0133 | 0.7972 ± 0.0210 | 0.8339 ± 0.0079 | <u>0.8558 ± 0.0177</u> | 0.8269 ± 0.0224 | 0.8317 ± 0.0219 |

Note: For each evaluation metric, the best-performing value is highlighted in **bold**, and the second-best is indicated with <u>underline</u> for ease of comparison.

**Table S4.** (continued)

| Endpoint | MoLFormer + Morgan FP | TF-IDF + D-MPNN | ChemBERTa + D-MPNN | MoLFormer + D-MPNN |
|---|---|---|---|---|
| CYP1A2 Inhibitor | 0.8572 ± 0.0021 | 0.8725 ± 0.0058 | **0.8856 ± 0.0060** | 0.8782 ± 0.0065 |
| CYP1A2 Substrate | <u>0.6084 ± 0.0338</u> | 0.5251 ± 0.0503 | 0.5427 ± 0.0547 | 0.5577 ± 0.0515 |
| CYP2C19 Inhibitor | 0.8235 ± 0.0145 | 0.8407 ± 0.0170 | **0.8588 ± 0.0065** | <u>0.8478 ± 0.0126</u> |
| CYP2C19 Substrate | <u>0.4908 ± 0.0438</u> | 0.3533 ± 0.0881 | 0.3524 ± 0.0628 | 0.3551 ± 0.0912 |
| CYP2C9 Inhibitor | 0.7860 ± 0.0120 | <u>0.8157 ± 0.0121</u> | **0.8258 ± 0.0047** | 0.8131 ± 0.0103 |
| CYP2C9 Substrate | 0.5987 ± 0.0468 | 0.5492 ± 0.0550 | <u>0.6180 ± 0.0636</u> | 0.5649 ± 0.0710 |
| CYP2D6 Inhibitor | 0.7042 ± 0.0102 | 0.7341 ± 0.0160 | <u>0.7531 ± 0.0046</u> | 0.7437 ± 0.0116 |
| CYP2D6 Substrate | 0.7071 ± 0.0244 | 0.6966 ± 0.0354 | 0.7086 ± 0.0286 | 0.6734 ± 0.0361 |
| CYP3A4 Inhibitor | 0.8160 ± 0.0053 | 0.8412 ± 0.0118 | <u>0.8455 ± 0.0063</u> | 0.8419 ± 0.0064 |
| CYP3A4 Substrate | 0.8052 ± 0.0134 | 0.7935 ± 0.0322 | 0.7786 ± 0.0188 | 0.7773 ± 0.0332 |
| OATP1B1 Inhibitor | 0.8280 ± 0.0104 | **0.8580 ± 0.0118** | <u>0.8577 ± 0.0096</u> | 0.8436 ± 0.0102 |
| OATP1B3 Inhibitor | 0.7374 ± 0.0239 | 0.7314 ± 0.0097 | 0.7427 ± 0.0148 | <u>0.7508 ± 0.0173</u> |
| OCT1 Inhibitor | 0.4565 ± 0.0289 | 0.4244 ± 0.0858 | 0.4432 ± 0.0628 | 0.4364 ± 0.0503 |
| OCT2 Inhibitor | 0.4770 ± 0.0281 | 0.4921 ± 0.0547 | 0.4880 ± 0.0265 | 0.5071 ± 0.0333 |
| UGTs Substrate | 0.6980 ± 0.0218 | 0.6703 ± 0.0460 | **0.7327 ± 0.0312** | 0.6819 ± 0.0463 |
| HLM Stability | 0.7507 ± 0.0186 | 0.7244 ± 0.0427 | **0.7665 ± 0.0322** | 0.7179 ± 0.0397 |
| RLM Stability | 0.8258 ± 0.0255 | 0.8380 ± 0.0125 | **0.8573 ± 0.0097** | 0.8436 ± 0.0170 |

Note: For each evaluation metric, the best-performing value is highlighted in **bold**, and the second-best is indicated with <u>underline</u> for ease of comparison.

**Table S5.** F1 scores for 1D+2D embedding combinations across classification endpoints

| Endpoint | 1D TF-IDF | 1D ChemBERTa | 1D MoLFormer | 2D Morgan FP | 2D D-MPNN | TF-IDF + Morgan FP | ChemBERTa + Morgan FP |
|---|---|---|---|---|---|---|---|
| CYP1A2 Inhibitor | 0.7548 ± 0.0104 | **0.7967 ± 0.0106** | 0.7611 ± 0.0112 | 0.7772 ± 0.0131 | 0.7951 ± 0.0129 | 0.7752 ± 0.0070 | 0.7757 ± 0.0059 |
| CYP1A2 Substrate | 0.3163 ± 0.0467 | 0.1305 ± 0.1281 | **0.4284 ± 0.0738** | 0.1765 ± 0.2163 | 0.3268 ± 0.1269 | 0.2741 ± 0.2263 | 0.3986 ± 0.1008 |
| CYP2C19 Inhibitor | 0.7404 ± 0.0085 | 0.7731 ± 0.0122 | 0.7492 ± 0.0096 | 0.7643 ± 0.0138 | <u>0.7854 ± 0.0105</u> | 0.7631 ± 0.0134 | 0.7685 ± 0.0106 |
| CYP2C19 Substrate | <u>0.1746 ± 0.1475</u> | 0.1083 ± 0.1783 | 0.1006 ± 0.1184 | 0.0519 ± 0.1037 | 0.0087 ± 0.0174 | 0.1084 ± 0.1660 | 0.1286 ± 0.1938 |
| CYP2C9 Inhibitor | 0.7092 ± 0.0107 | 0.7355 ± 0.0045 | 0.7157 ± 0.0123 | 0.7073 ± 0.0100 | 0.7525 ± 0.0098 | 0.7173 ± 0.0177 | 0.7241 ± 0.0090 |
| CYP2C9 Substrate | 0.4321 ± 0.0339 | 0.1165 ± 0.1652 | 0.4097 ± 0.2119 | 0.3277 ± 0.2079 | 0.2129 ± 0.1855 | 0.3659 ± 0.2013 | 0.4137 ± 0.1597 |
| CYP2D6 Inhibitor | 0.6030 ± 0.0163 | 0.6400 ± 0.0113 | 0.5892 ± 0.0170 | 0.5916 ± 0.0260 | <u>0.6707 ± 0.0100</u> | 0.6120 ± 0.0123 | 0.6235 ± 0.0069 |
| CYP2D6 Substrate | 0.5738 ± 0.0285 | 0.5981 ± 0.0895 | 0.5679 ± 0.0433 | 0.6396 ± 0.0405 | 0.5409 ± 0.0343 | <u>0.6747 ± 0.0666</u> | **0.6899 ± 0.0121** |
| CYP3A4 Inhibitor | 0.7194 ± 0.0084 | 0.7493 ± 0.0055 | 0.7305 ± 0.0073 | 0.7435 ± 0.0049 | **0.7741 ± 0.0082** | 0.7351 ± 0.0068 | 0.7443 ± 0.0162 |
| CYP3A4 Substrate | 0.7539 ± 0.0210 | 0.7606 ± 0.0281 | 0.7267 ± 0.0264 | **0.7830 ± 0.0159** | 0.7306 ± 0.0135 | <u>0.7635 ± 0.0072</u> | 0.7569 ± 0.0275 |
| OATP1B1 Inhibitor | 0.8081 ± 0.0169 | 0.8243 ± 0.0170 | **0.8276 ± 0.0218** | 0.8179 ± 0.0139 | 0.8074 ± 0.0177 | 0.8209 ± 0.0102 | 0.8245 ± 0.0026 |
| OATP1B3 Inhibitor | 0.7082 ± 0.0154 | 0.6970 ± 0.0297 | 0.7099 ± 0.0216 | 0.6785 ± 0.0450 | 0.7348 ± 0.0143 | 0.6921 ± 0.0363 | 0.7173 ± 0.0188 |
| OCT1 Inhibitor | 0.2148 ± 0.1453 | 0.0426 ± 0.0851 | <u>0.2434 ± 0.1644</u> | 0.1490 ± 0.1551 | 0.2142 ± 0.1613 | 0.1780 ± 0.1624 | 0.2017 ± 0.1640 |
| OCT2 Inhibitor | 0.2127 ± 0.1078 | 0.2268 ± 0.0470 | 0.3529 ± 0.0921 | 0.1580 ± 0.1607 | **0.4594 ± 0.0408** | 0.2851 ± 0.1470 | 0.3284 ± 0.0933 |
| UGTs Substrate | 0.5231 ± 0.1936 | <u>0.6788 ± 0.0349</u> | 0.5752 ± 0.0622 | 0.4489 ± 0.1511 | 0.6057 ± 0.0471 | 0.5861 ± 0.0711 | 0.6416 ± 0.0771 |
| HLM Stability | 0.4895 ± 0.2564 | <u>0.6960 ± 0.0122</u> | 0.6476 ± 0.0097 | 0.6792 ± 0.0082 | 0.6623 ± 0.0241 | 0.6949 ± 0.0201 | 0.6984 ± 0.0312 |
| RLM Stability | 0.7665 ± 0.0196 | 0.7642 ± 0.0110 | 0.7388 ± 0.0209 | 0.7692 ± 0.0092 | <u>0.7793 ± 0.0183</u> | 0.7611 ± 0.0150 | 0.7581 ± 0.0104 |

Note: For each evaluation metric, the best-performing value is highlighted in **bold**, and the second-best is indicated with <u>underline</u> for ease of comparison.

**Table S5.** (continued)

| Endpoint | MoLFormer + Morgan FP | TF-IDF + D-MPNN | ChemBERTa + D-MPNN | MoLFormer + D-MPNN |
|---|---|---|---|---|
| CYP1A2 Inhibitor | 0.7722 ± 0.0067 | 0.7835 ± 0.0118 | <u>0.7960 ± 0.0076</u> | 0.7871 ± 0.0079 |
| CYP1A2 Substrate | 0.4923 ± 0.0894 | 0.2925 ± 0.1058 | 0.3820 ± 0.0902 | <u>0.4207 ± 0.1459</u> |
| CYP2C19 Inhibitor | 0.7602 ± 0.0104 | 0.7790 ± 0.0114 | **0.7881 ± 0.0063** | 0.7820 ± 0.0106 |
| CYP2C19 Substrate | **0.2107 ± 0.0356** | 0.0747 ± 0.1494 | 0.0658 ± 0.0827 | 0.0742 ± 0.0943 |
| CYP2C9 Inhibitor | 0.7191 ± 0.0129 | 0.7554 ± 0.0137 | **0.7609 ± 0.0092** | <u>0.7542 ± 0.0111</u> |
| CYP2C9 Substrate | <u>0.4743 ± 0.1336</u> | 0.4293 ± 0.2186 | **0.5717 ± 0.0898** | 0.4290 ± 0.1564 |
| CYP2D6 Inhibitor | 0.6256 ± 0.0087 | 0.6571 ± 0.0110 | **0.6729 ± 0.0046** | 0.6671 ± 0.0099 |
| CYP2D6 Substrate | 0.6731 ± 0.0274 | 0.6316 ± 0.0546 | 0.6406 ± 0.0276 | 0.5953 ± 0.0416 |
| CYP3A4 Inhibitor | 0.7432 ± 0.0133 | 0.7664 ± 0.0073 | <u>0.7735 ± 0.0061</u> | 0.7670 ± 0.0060 |
| CYP3A4 Substrate | 0.7578 ± 0.0181 | 0.7282 ± 0.0084 | 0.7285 ± 0.0074 | 0.7352 ± 0.0307 |
| OATP1B1 Inhibitor | 0.8272 ± 0.0033 | <u>0.8248 ± 0.0083</u> | 0.8246 ± 0.0043 | 0.8239 ± 0.0075 |
| OATP1B3 Inhibitor | 0.7210 ± 0.0153 | <u>0.7352 ± 0.0129</u> | 0.7299 ± 0.0150 | **0.7362 ± 0.0204** |
| OCT1 Inhibitor | 0.2328 ± 0.1796 | 0.2198 ± 0.1145 | **0.2984 ± 0.0923** | 0.2150 ± 0.1722 |
| OCT2 Inhibitor | 0.2697 ± 0.1644 | 0.3767 ± 0.0399 | 0.4408 ± 0.0529 | <u>0.4458 ± 0.0345</u> |
| UGTs Substrate | 0.6685 ± 0.0245 | 0.6260 ± 0.0576 | **0.6957 ± 0.0396** | 0.6292 ± 0.0840 |
| HLM Stability | 0.6960 ± 0.0166 | 0.6484 ± 0.0442 | **0.6987 ± 0.0261** | 0.6515 ± 0.0415 |
| RLM Stability | 0.7496 ± 0.0166 | 0.7777 ± 0.0121 | **0.7817 ± 0.0109** | 0.7769 ± 0.0172 |

Note: For each evaluation metric, the best-performing value is highlighted in **bold**, and the second-best is indicated with <u>underline</u> for ease of comparison.

**Table S6.** Performance comparison for 1D+2D embedding combinations across regression endpoints

| Embedding | Clearance | | | Half-life | | |
|---|---|---|---|---|---|---|
| | RMSE | MAE | $R^2$ | RMSE | MAE | $R^2$ |
| 1D TF-IDF | 0.7647 ± 0.0418 | 0.5135 ± 0.0193 | 0.3794 ± 0.0610 | 0.6924 ± 0.0384 | 0.4958 ± 0.0132 | 0.1935 ± 0.0532 |
| 1D ChemBERTa | 0.7681 ± 0.0434 | 0.5165 ± 0.0314 | 0.3751 ± 0.0468 | 0.7017 ± 0.0459 | 0.5002 ± 0.0232 | 0.1738 ± 0.0384 |
| 1D MoLFormer | 0.7868 ± 0.0276 | 0.5355 ± 0.0199 | 0.3427 ± 0.0562 | 0.7201 ± 0.0378 | 0.5190 ± 0.0136 | 0.1272 ± 0.0596 |
| 2D Morgan FP | 0.7227 ± 0.0197 | 0.4945 ± 0.0157 | 0.4452 ± 0.0463 | 0.6908 ± 0.0438 | 0.4953 ± 0.0225 | 0.1971 ± 0.0680 |
| 2D D-MPNN | 0.8002 ± 0.0263 | 0.5434 ± 0.0098 | 0.3219 ± 0.0249 | 0.6922 ± 0.0532 | 0.4964 ± 0.0218 | 0.1936 ± 0.0830 |
| TF-IDF + Morgan FP | **0.7119 ± 0.0320** | **0.4826 ± 0.0131** | <u>0.4592 ± 0.0724</u> | 0.6872 ± 0.0385 | 0.4973 ± 0.0187 | 0.2051 ± 0.0640 |
| ChemBERTa + Morgan FP | <u>0.7120 ± 0.0267</u> | <u>0.4853 ± 0.0258</u> | **0.4609 ± 0.0548** | 0.6911 ± 0.0379 | 0.4954 ± 0.0183 | 0.1943 ± 0.0828 |
| MoLFormer + Morgan FP | 0.7186 ± 0.0304 | 0.4865 ± 0.0229 | 0.4505 ± 0.0623 | 0.6998 ± 0.0469 | 0.5038 ± 0.0264 | 0.1744 ± 0.0913 |
| TF-IDF + D-MPNN | 0.7363 ± 0.0402 | 0.4984 ± 0.0165 | 0.4260 ± 0.0385 | <u>0.6821 ± 0.0448</u> | <u>0.4886 ± 0.0157</u> | 0.2170 ± 0.0684 |
| ChemBERTa + D-MPNN | 0.7774 ± 0.0093 | 0.5221 ± 0.0181 | 0.3579 ± 0.0499 | **0.6768 ± 0.0517** | **0.4814 ± 0.0219** | **0.2303 ± 0.0672** |
| MoLFormer + D-MPNN | 0.7689 ± 0.0117 | 0.5259 ± 0.0125 | 0.3725 ± 0.0387 | 0.6822 ± 0.0487 | 0.4920 ± 0.0168 | <u>0.2179 ± 0.0630</u> |

Note: For each evaluation metric, the best-performing value is highlighted in **bold**, and the second-best is indicated with <u>underline</u> for ease of comparison.

**Table S7.** AUROC scores for fusion strategies across classification endpoints

| Endpoint | Concatenation | Linear Attention | CNN-based Attention | Multi-view Attention |
|---|---|---|---|---|
| CYP1A2 Inhibitor | <u>0.8939 ± 0.0054</u> | 0.8926 ± 0.0066 | 0.8930 ± 0.0063 | **0.9010 ± 0.0067** |
| CYP1A2 Substrate | <u>0.8414 ± 0.0195</u> | 0.8287 ± 0.0226 | 0.8153 ± 0.0435 | **0.8575 ± 0.0090** |
| CYP2C19 Inhibitor | <u>0.8923 ± 0.0048</u> | 0.8913 ± 0.0045 | 0.8910 ± 0.0044 | **0.8983 ± 0.0067** |
| CYP2C19 Substrate | 0.7832 ± 0.0495 | 0.7449 ± 0.0688 | **0.8082 ± 0.0174** | <u>0.8011 ± 0.0062</u> |
| CYP2C9 Inhibitor | <u>0.8823 ± 0.0038</u> | 0.8808 ± 0.0049 | 0.8818 ± 0.0045 | **0.8854 ± 0.0061** |
| CYP2C9 Substrate | <u>0.8619 ± 0.0141</u> | 0.8180 ± 0.0731 | 0.8531 ± 0.0200 | **0.8705 ± 0.0166** |
| CYP2D6 Inhibitor | **0.8635 ± 0.0023** | 0.8626 ± 0.0034 | <u>0.8632 ± 0.0043</u> | 0.8631 ± 0.0053 |
| CYP2D6 Substrate | <u>0.8736 ± 0.0055</u> | 0.8696 ± 0.0154 | 0.8628 ± 0.0168 | **0.8994 ± 0.0103** |
| CYP3A4 Inhibitor | 0.8730 ± 0.0049 | 0.8727 ± 0.0050 | <u>0.8733 ± 0.0061</u> | **0.8757 ± 0.0051** |
| CYP3A4 Substrate | 0.8538 ± 0.0103 | <u>0.8562 ± 0.0249</u> | 0.8503 ± 0.0253 | **0.8809 ± 0.0153** |
| OATP1B1 Inhibitor | 0.7325 ± 0.0155 | 0.7326 ± 0.0197 | <u>0.7364 ± 0.0133</u> | **0.7419 ± 0.0131** |
| OATP1B3 Inhibitor | **0.7168 ± 0.0133** | 0.7154 ± 0.0165 | 0.7125 ± 0.0179 | <u>0.7158 ± 0.0145</u> |
| OCT1 Inhibitor | **0.7775 ± 0.0370** | 0.7674 ± 0.0296 | <u>0.7762 ± 0.0314</u> | 0.7760 ± 0.0322 |
| OCT2 Inhibitor | 0.8165 ± 0.0191 | 0.8044 ± 0.0316 | <u>0.8205 ± 0.0120</u> | **0.8271 ± 0.0213** |
| UGTs Substrate | **0.8873 ± 0.0143** | 0.8813 ± 0.0163 | <u>0.8850 ± 0.0156</u> | 0.8797 ± 0.0158 |
| HLM Stability | <u>0.8574 ± 0.0156</u> | 0.8512 ± 0.0118 | 0.8517 ± 0.0264 | **0.8633 ± 0.0103** |
| RLM Stability | **0.8224 ± 0.0096** | 0.8170 ± 0.0085 | 0.8165 ± 0.0087 | <u>0.8176 ± 0.0104</u> |

Note: For each evaluation metric, the best-performing value is highlighted in **bold**, and the second-best is indicated with <u>underline</u> for ease of comparison.

**Table S8.** AUPRC scores for fusion strategies across classification endpoints

| Endpoint | Concatenation | Linear Attention | CNN-based Attention | Multi-view Attention |
|---|---|---|---|---|
| CYP1A2 Inhibitor | <u>0.8856 ± 0.0060</u> | 0.8841 ± 0.0075 | 0.8827 ± 0.0070 | **0.8903 ± 0.0083** |
| CYP1A2 Substrate | <u>0.5427 ± 0.0547</u> | 0.5296 ± 0.0748 | 0.4684 ± 0.0966 | **0.6059 ± 0.0165** |
| CYP2C19 Inhibitor | <u>0.8588 ± 0.0065</u> | 0.8580 ± 0.0091 | 0.8584 ± 0.0088 | **0.8712 ± 0.0071** |
| CYP2C19 Substrate | 0.3524 ± 0.0628 | 0.3034 ± 0.0868 | <u>0.3782 ± 0.0464</u> | **0.4353 ± 0.0701** |
| CYP2C9 Inhibitor | 0.8258 ± 0.0047 | <u>0.8261 ± 0.0042</u> | 0.8251 ± 0.0037 | **0.8361 ± 0.0057** |
| CYP2C9 Substrate | <u>0.6180 ± 0.0636</u> | 0.5044 ± 0.1312 | 0.5857 ± 0.0909 | **0.6498 ± 0.0606** |
| CYP2D6 Inhibitor | **0.7531 ± 0.0046** | 0.7508 ± 0.0114 | 0.7519 ± 0.0102 | <u>0.7526 ± 0.0122</u> |
| CYP2D6 Substrate | <u>0.7086 ± 0.0286</u> | 0.6729 ± 0.0536 | 0.6687 ± 0.0533 | **0.7611 ± 0.0183** |
| CYP3A4 Inhibitor | 0.8455 ± 0.0063 | 0.8452 ± 0.0062 | <u>0.8470 ± 0.0073</u> | **0.8499 ± 0.0080** |
| CYP3A4 Substrate | <u>0.7786 ± 0.0188</u> | 0.7691 ± 0.0346 | 0.7649 ± 0.0423 | **0.8312 ± 0.0274** |
| OATP1B1 Inhibitor | 0.8577 ± 0.0096 | 0.8557 ± 0.0063 | <u>0.8623 ± 0.0055</u> | **0.8629 ± 0.0095** |
| OATP1B3 Inhibitor | 0.7427 ± 0.0148 | <u>0.7439 ± 0.0240</u> | 0.7419 ± 0.0207 | **0.7492 ± 0.0238** |
| OCT1 Inhibitor | <u>0.4432 ± 0.0628</u> | 0.4367 ± 0.0509 | 0.4350 ± 0.0618 | **0.4738 ± 0.0626** |
| OCT2 Inhibitor | 0.4880 ± 0.0265 | 0.4801 ± 0.0397 | <u>0.4966 ± 0.0361</u> | **0.5242 ± 0.0359** |
| UGTs Substrate | <u>0.7327 ± 0.0312</u> | 0.7105 ± 0.0248 | 0.7243 ± 0.0322 | **0.7398 ± 0.0465** |
| HLM Stability | <u>0.7665 ± 0.0322</u> | 0.7533 ± 0.0274 | 0.7573 ± 0.0502 | **0.7864 ± 0.0167** |
| RLM Stability | <u>0.8573 ± 0.0097</u> | 0.8524 ± 0.0136 | 0.8503 ± 0.0092 | **0.8591 ± 0.0070** |

Note: For each evaluation metric, the best-performing value is highlighted in **bold**, and the second-best is indicated with <u>underline</u> for ease of comparison.

**Table S9.** F1 scores for fusion strategies across classification endpoints

| Endpoint | Concatenation | Linear Attention | CNN-based Attention | Multi-view Attention |
|---|---|---|---|---|
| CYP1A2 Inhibitor | <u>0.7960 ± 0.0076</u> | 0.7954 ± 0.0143 | 0.7904 ± 0.0089 | **0.8132 ± 0.0094** |
| CYP1A2 Substrate | <u>0.3820 ± 0.0902</u> | 0.2976 ± 0.2055 | 0.2301 ± 0.1551 | **0.5706 ± 0.0312** |
| CYP2C19 Inhibitor | <u>0.7881 ± 0.0063</u> | 0.7875 ± 0.0109 | 0.7830 ± 0.0079 | **0.8024 ± 0.0101** |
| CYP2C19 Substrate | 0.0658 ± 0.0827 | 0.0577 ± 0.0991 | <u>0.1292 ± 0.1194</u> | **0.4510 ± 0.0578** |
| CYP2C9 Inhibitor | <u>0.7609 ± 0.0092</u> | 0.7604 ± 0.0059 | 0.7569 ± 0.0130 | **0.7708 ± 0.0109** |
| CYP2C9 Substrate | <u>0.5717 ± 0.0898</u> | 0.3891 ± 0.2157 | 0.4441 ± 0.2176 | **0.6074 ± 0.0516** |
| CYP2D6 Inhibitor | 0.6729 ± 0.0046 | <u>0.6736 ± 0.0100</u> | 0.6637 ± 0.0127 | **0.6905 ± 0.0098** |
| CYP2D6 Substrate | <u>0.6406 ± 0.0276</u> | 0.5339 ± 0.2710 | 0.6142 ± 0.0638 | **0.7104 ± 0.0435** |
| CYP3A4 Inhibitor | 0.7735 ± 0.0061 | <u>0.7747 ± 0.0059</u> | 0.7716 ± 0.0065 | **0.7827 ± 0.0043** |
| CYP3A4 Substrate | <u>0.7285 ± 0.0074</u> | 0.7220 ± 0.0627 | 0.6722 ± 0.1300 | **0.7895 ± 0.0169** |
| OATP1B1 Inhibitor | <u>0.8246 ± 0.0043</u> | 0.8231 ± 0.0074 | 0.8242 ± 0.0072 | **0.8338 ± 0.0046** |
| OATP1B3 Inhibitor | 0.7299 ± 0.0150 | <u>0.7337 ± 0.0213</u> | 0.7297 ± 0.0245 | **0.7557 ± 0.0141** |
| OCT1 Inhibitor | <u>0.2984 ± 0.0923</u> | 0.1496 ± 0.1437 | 0.1860 ± 0.1716 | **0.5222 ± 0.0238** |
| OCT2 Inhibitor | <u>0.4408 ± 0.0529</u> | 0.3493 ± 0.1611 | 0.4091 ± 0.1206 | **0.5466 ± 0.0419** |
| UGTs Substrate | 0.6957 ± 0.0396 | <u>0.6962 ± 0.0335</u> | 0.6936 ± 0.0304 | **0.6969 ± 0.0234** |
| HLM Stability | <u>0.6987 ± 0.0261</u> | 0.6972 ± 0.0176 | 0.6674 ± 0.0454 | **0.7104 ± 0.0233** |
| RLM Stability | 0.7817 ± 0.0109 | <u>0.7826 ± 0.0110</u> | 0.7787 ± 0.0088 | **0.7883 ± 0.0079** |

Note: For each evaluation metric, the best-performing value is highlighted in **bold**, and the second-best is indicated with <u>underline</u> for ease of comparison.

**Table S10.** Performance comparison for fusion strategies across regression endpoints

| Embedding | Clearance | | | Half-life | | |
|---|---|---|---|---|---|---|
| | **RMSE** | **MAE** | **R2** | **RMSE** | **MAE** | **R2** |
| Concatenation | 0.7774 ± 0.0093 | 0.5221 ± 0.0181 | 0.3579 ± 0.0499 | 0.6768 ± 0.0517 | 0.4814 ± 0.0219 | 0.2303 ± 0.0672 |
| Linear Attention | <u>0.7564 ± 0.0192</u> | <u>0.5056 ± 0.0138</u> | <u>0.3918 ± 0.0529</u> | <u>0.6731 ± 0.0431</u> | <u>0.4799 ± 0.0176</u> | <u>0.2377 ± 0.0632</u> |
| CNN-based Attention | 0.7657 ± 0.0511 | 0.5101 ± 0.0292 | 0.3786 ± 0.0589 | 0.6750 ± 0.0445 | 0.4822 ± 0.0154 | 0.2330 ± 0.0716 |
| Multi-view Attention | **0.6856 ± 0.0448** | **0.4469 ± 0.0163** | **0.5000 ± 0.0636** | **0.6683 ± 0.0547** | **0.4735 ± 0.0242** | **0.2494 ± 0.0740** |

Note: For each evaluation metric, the best-performing value is highlighted in **bold**, and the second-best is indicated with <u>underline</u> for ease of comparison.

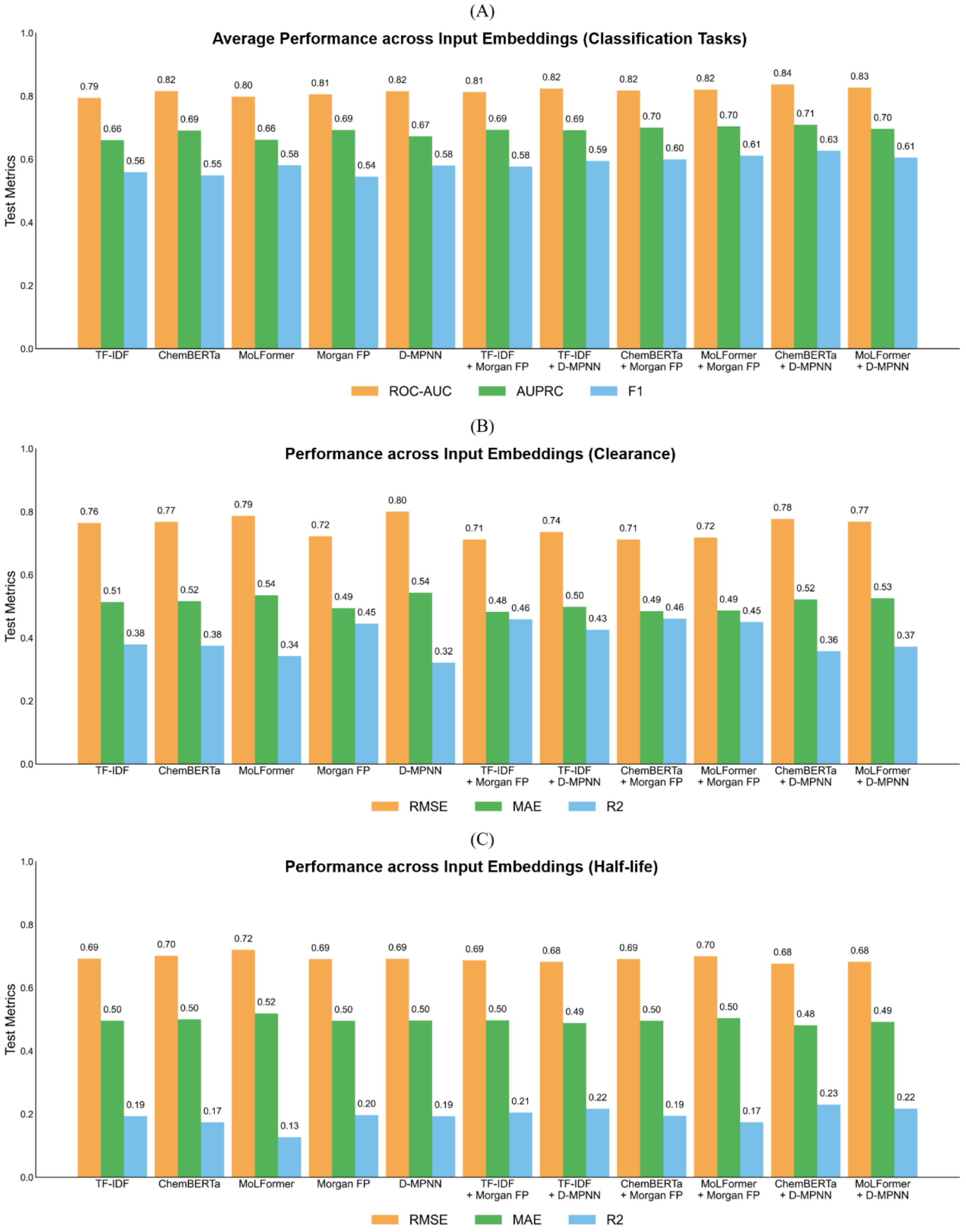


**Figure S1.** Performance for input single embeddings and 1D+2D embedding combinations across ME endpoints. (A) Average performance across input embeddings for classification tasks. (B) Performance across input embeddings for clearance. (C) Performance across input embeddings for half-life.
